# Haplotype-phased and chromosome-level genome assembly of *Puccinia polysora*, a giga-scale fungal pathogen causing southern corn rust

**DOI:** 10.1101/2022.05.18.492400

**Authors:** Junmin Liang, Yuanjie Li, Peter N. Dodds, Melania Figueroa, Jana Sperschneider, Shiling Han, Clement K.M. Tsui, Keyu Zhang, Leifu Li, Zhanhong Ma, Lei Cai

**Author notes:** Correspondence: Lei Cai, State Key Laboratory of Mycology, Institute of Microbiology, Chinese Academy of Sciences, Beijing, 100101, China.

## Abstract

Rust fungi are characterized by large genomes with high repeat content and have two haploid nuclei in most life stages, which makes achieving high-quality genome assemblies challenging. Here, we described a pipeline using HiFi reads and Hi-C data to assemble a gigabase-sized fungal pathogen, *Puccinia polysora* f.sp. *zeae*, to haplotype-phased and chromosome-scale. The final assembled genome is 1.71 Gbp, with ~850 Mbp and 18 chromosomes in each haplotype, being currently one of the two giga-scale fungi assembled to chromosome level. Transcript-based annotation identified 47,512 genes for dikaryotic genome with a similar number for each haplotype. A high level of interhaplotype variation was found with 10% haplotype-specific BUSCO genes, 5.8 SNPs/kbp and structural variation accounting for 3% of the genome size. The *P. polysora* genome displayed over 85% repeat contents, with genome-size expansion and copy number increasing of species-specific orthogroups. Interestingly, these features did not affect overall synteny with other *Puccinia* species having smaller genomes. Fine-time-point transcriptomics revealed seven clusters of co-expressed secreted proteins that are conserved between two haplotypes. The fact that candidate effectors interspersed with all genes indicated the absence of a “two-speed genome” evolution in *P. polysora*. Genome resequencing of 79 additional isolates revealed a clonal population structure of *P. polysora* in China with low geographic differentiation. Nevertheless, a minor population differentiated from the major population by having mutations on secreted proteins including *AvrRppC*, indicating the ongoing virulence to evade recognition by *RppC*, a major resistance gene in Chinese corn cultivars. The high-quality assembly provides valuable genomic resources for future studies on disease management and the evolution of *P. polysora*.

## 1 INTRODUCTION

Rust fungi (Pucciniales) constitute one of the largest orders in the kingdom of fungi, with more than 8,000 species grouped into 18 families and approximately 170 accepted genera (Zhao et al. 2020, 2021; Aime and McTaggart 2021). Rust fungi are obligate biotrophs, which are usually recalcitrant to in vitro culturing and are host-specific to particular species, genera, or families of vascular plants. Their life cycles are diverse and complex, including up to five types of spores (spermatia, aeciospores, urediniospores, teliospores, and basidiospores) (Zhao et al. 2021). Spores of the most abundant life stage are dikaryon, which contains two physically separated and genetically different haploid nuclei. The Pucciniales contains species with the largest known genomes, with an average haplotype genome size of 380 Mbp estimated by flow cytometry, far larger than that of all fungi (37.7 Mbp) (Tavares et al. 2014). Based on flow cytometry, the largest known rust genome (*Uromyces bidentis*) was estimated at up to 2.4 Gbp (Ramos et al. 2015). The repeat contents of rust genomes range from 30% to 91% (Amie et al. 2017; Tobias et al. 2021). Their dikaryotic nature and highly repetitive sequences pose substantial challenges for high-quality genome assembly as well as an in-depth understanding of evolution within the rust group.

In addition to the above intriguing biological features, rust fungi have also received wide attention for causing major diseases on agricultural and forest crops worldwide. The most devastating rust pathogen include *Puccinia striiformis* f.sp. *tritici* (*Pst*), *P. graminis* f.sp. *tritici* (*Pgf*) and *P. triticina* f.sp. *tritici* (*Ptt*), causing three of the world’s most serious wheat rust diseases (Kolmer 2005); *Puccinia sorghi* f.sp. *zeae* (*Psz*) and *P. polysora* f.sp. *zeae* (*Ppz*), also seriously threaten global crop production by causing common and southern corn rust of maize (Crouch and Szabo 2011; Ramirez-Cabral et al. 2017); and *Melampsora larici-populina* and *Austropuccinia psidii*, two notorious forest pathogens leading to poplar rust and myrtle rust, respectively (Pinon and Frey 1997; Winzer et al. 2019). Effective strategies for management of rust disease rely heavily on host resistance breeding programs. In host-rust interactions, rust fungi deliver specific effectors (avirulence genes, *Avr*) into host cells, which can be recognized by resistance proteins from resistant plants and trigger defense responses (Dodds and Rathjen 2010). However, rust pathogens can acquire mutations in the *Avr* gene to avoid this recognition and facilitate the pathogen infection (Cui et al. 2015). For instance, the Ug99 race (TTKSK) of *P. graminis* evolved virulence to the widely deployed *Sr31* resistance gene in wheat has become a big threat to global wheat production (Singh et al. 2011). Therefore, understanding the molecular mechanism of *Avr-R*, or gene-for-gene relationship, is critical for durable disease control. Given their biotrophic lifestyle, the genetic transformation of rust fungi is difficult and challenging. Nevertheless, genome sequencing data of rust species has enabled the prediction of candidate effectors which facilitate the identification of *Avr* genes (Figueroa et al. 2016; Anderson et al. 2016; Maia et al. 2017; Salcedo et al. 2017; Chen et al. 2017; Miller et al. 2018; 2020; Upadhyaya et al 2021).

Early rust genome assemblies were haploid representations and different homologous haplotypes were collapsed into a consensus assembly, which did not fully capture sequences from both nuclei. Using single-molecule real-time (SMRT) sequencing technology and haplotype resolution software, FALCON-Unzip (Chin et al. 2016), allowed higher contiguity and partially haplotype-phased genomes of a few *Puccinia* species (Miller et al. 2018; Schwessinger et al. 2018, Wu et al. 2021). Other softwares have been developed to obtain sub-assemblies from diploid assemblies, such as HaploMerger2 (Huang et al. 2017) and Purge_Haplotigs (Roach et al. 2018), but many duplicate contigs cannot be correctly switched leading to mis-joins in the final sub-assemblies. Recently, several softwares have been developed to phase two haplotypes from the diploid genome using Hi-C data, such as TrioCanu (Koren et al. 2018), NuclearPhaser (Duan et al. 2022) and FALCON-Phase (Kronenberg et al. 2022). Three *Puccinia* species, causing wheat or oat rust have been assembled to a fully-phased and chromosome level using NuclearPhaser (Li et al. 2019; Duan et al. 2022; Henningsen et al. 2022). Unlike above rusts, which have been studied extensively, the genome information of corn rust pathogens is largely unknown. Since 2012, global maize production has exceeded that of wheat and rice, becoming the food crop with the highest world production yield (FAO, https://www.fao.org/faostat/en/#data/QCL), meanwhile, the distribution of the two major rust diseases (common corn rust and southern corn rust) in corn is projected to expand to temperate regions with increasing global temperatures (Ramirez-Cabral et al. 2017), which poses a greater threat to global food safety. A draft genome sequence of common corn rust (*P. sorghi*) was released in 2016, although it was highly fragmented (Rochi et al. 2016). However, genomic information for southern corn rust pathogens has not been reported, which seriously hampers avirulence gene identification and breeding of disease resistance. The genome sizes of recent phased *Puccinia* species range from ~170 Mbp to 250 Mbp (Li et al. 2019; Miller et al. 2018; Wu et al. 2021), whereas the genome of *P. polysora* was estimated over gigabase in our preliminary test (Figure S1), which poses additional challenges for genome assembly.

In this study, we assembled the genome of *P. polysora* to haplotype-resolved and chromosome-scale levels using a modified haplotype-phasing assembly pipeline. To our knowledge, the current study and the report on the myrtle rust pathogen, *Austropuccinia psidii* (Edwards et al. 2022) provide the first giga-scale fungal genome ever assembled to complete-chromosome level. The *P. polysora* genome was annotated by fine-time-point gene expression data from germinated spores and infected issues, which supplied robust data for coexpression analyses of secreted proteins and prediction of candidate secreted effectors. Also, using the haplotype-phasing genome as a reference, we investigated the genetic and population structure of *P. polysora* based on genome sequencing of 79 additional samples from north, south and central parts of China. The high-quality genome information represents a valuable resource for better understanding genome evolution, host adaption, and genes involved in host-pathogen interactions.

## 2 MATERIALS AND METHODS

### 2.1 Fungal isolates and plant inoculation

The isolate, GD1913, collected from Guangdong province in the 2019 annual rust survey, was chosen for reference genome sequencing. Based on a preliminary inoculation experiment, GD1913 represented the most frequent virulence type. Additional 79 isolates, collected from the south, central, and north China were selected and used for genome resequencing (Table S1). All isolates were purified by selecting a single pustule from infected leaves and amplified by 2–3 rounds of infection on highly susceptible variety Zhengdan 958. Corn seeds immersed in Chlormequat chloride (1.5 g/L) were sown in 10-cm wide square pots 10 days before inoculation. When 10-cm tall, seedling growth was reduced by adding 15 mL maleic hydrazide acid (1.5 g/L) per pot. Urediniospores were amplified by spraying on 10-day-old corn seedlings. The inoculated seedlings were incubated with dew at 27°C in the dark for 24h before transferring to a climate-controlled chamber (12h, 25°C dark period and a 12h, 27°C light period). To prevent airborne contamination, 75% ethyl alcohol was sprayed on the area after each inoculation and each pot was protected by a cellophane bag. Once the pots were heavily infected (ca. 15–20 days), spores were harvested to clean cellophane by scraping them with sterile needles. For long-term storage, fresh spores were dried (10% relative humidity) in a desiccator for 1 day at 4°C and then maintained at −80°C or in liquid nitrogen.

### 2.2 DNA extraction, library preparation, and sequencing

About 500 mg of fresh urediniospores were ground for 4-5 batches in liquid nitrogen and genomic DNA was extracted using the lysis buffer and cetyltrimethylammonium bromide (CTAB) method (Justesen et al. 2002). The integrity of genomic DNA was assessed by Agilent 4200 Bioanalyzer (Agilent Technologies, Palo Alto, California, USA). The average insert size of 15 kb PacBio library was concentrated with AMPure PB magnetic beads (Pacific Biosciences, California, USA) following the manufacturer’s instruction. Sequencing of high fidelity (HiFi) long reads was carried out by the Pacific Bioscience Sequel II platform. For genome resequencing, ~10 mg fresh urediniospores were placed in 2 mL screw-gap tubes filled with Lysing Matrix C and ground twice in MP FastPrep-24 TM 5G (Mp Biomedicals, USA) with a speed setting of 4 for 20 seconds. The paired-end library with 150 bp was prepared and sequenced using Illumina NovaSeq 6000. All sequencing was performed at Annoroad Gene Technology Co., Ltd, (Beijing, China).

To generate a chromosome-level and haplotype phased assembly of *P. polysora* genome, a Hi-C library was generated following in situ ligation protocols. In brief, fresh urediniospores (~ 100 mg) of GD1913 were used for crosslinking reaction by 2% formaldehyde at room temperature for 15 min. After Glycine quenching, the supernatant was removed and spores were then ground with liquid nitrogen for DNA extraction. The purified DNA was digested with *Mbo* I restriction enzyme (New England Biolabs Inc. Beijing, China) and was labeled by incubating with Biotin-14-dATP (Thermo Fisher Scientific, Massachusetts, USA) and then ligated by T4

DNA Ligase (Thermo Fisher Scientific, Massachusetts, USA). After incubating overnight to reverse crosslinks, the ligated DNA was sheared into ~350 bp fragments. Finally, the Hi-C library was quantified by Bioanalyzer and sequenced on the Illumina NovaSeq 6000 platform using paired-end 150 cycles.

### 2.3 *De novo* assembly, haplotype phasing and Hi-C scaffolding

HiFi reads were assembled by Canu 2.1.1 with -pacbio-hifi (Nurk et al. 2020). The coverage depth was calculated by genomeCoverageBed in BEDtools (v2.29.2) (Quinlan and Hall 2010) and small contigs (< 20 kbp) with low coverage (< 2×) were excluded from the further assembly. The remaining contigs were examined by BLASTN search (v2.7.2) against the NCBI nt/nr database (downloaded on May 20, 2021) with E-value set as 1e-10. To move the contaminants e.g. plant rDNA and chloroplast sequences, we utilized a ContaminantScreening pipeline (https://github.com/JanaSperschneider/GenomeAssemblyTools/tree/master/ContaminantScreening). Contigs of high coverage and low GC content were defined as potential mitochondrial contigs, and they were extracted from the assembled output from Canu and searched against the mitochondrial database from NCBI (https://ftp.ncbi.nlm.nih.gov/refseq/release/mitochondrion/). The contigs (63 contigs in Table S2) with perfect matches to mitochondria were also excluded and they were assembled separately by Canu v2.1.1 to obtain mtDNA. The assembled sequences were searched against itself database to test the ring formation.

To obtain a haplotype-phased assembly, HaploMerger2 (Huang et al. 2017), a tool to rebuild both haploid sub-assemblies from the high-heterozygosity diploid genome, was used. The heterozygosity evaluated by Jellyfish v2.1.3 (Marçais and Kingsford 2011) and Genomescope.R (Vurture et al. 2017) was 1.08% leading to the identity setting as 95% (Huang et al. 2017). To better distinguish allelic and non-allelic combinations, the scoring scheme for alignments was recalculated by a Perl script (lastz_D_Wrapper.pl) that came with HaploMerger2. The top 22 (in length) contigs accounting for ~10% length of all contigs were assigned to part1.fasta and others were in part2.fasta. Mis-joins were detected by three rounds and all breaks were manually checked combined with Hi-C validation. Based on all-vs-all alignment (hm.new_scaffolds), contigs were assigned separately to haplotype A and haplotype B. For scaffolding, the raw Hi-C data were trimmed by removing adapters and low-quality bases, and then two-paired reads were separately mapped to two haplotypes independently using BWA-MEM v0.7.8 (Li and Durbin 2010). To move experimental artifacts, the alignments went through the mapping workflow from the Arima Genomics pipeline (https://github.com/ArimaGenomics/mapping_pipeline/blob/master/01_mapping_arima.sh). Then SALSA v2.2 (Ghurye et al. 2017) was run to cluster initial contigs into groups. SALSA scaffolding was performed independently on haplotype A and B contigs. Hi-C contact matrices for each scaffold were calculated by HiC-Pro (Servant et al. 2015) and contact maps are visualized by HiCPlotter (Akdemir and Chin 2015). Contig reversal and breaking of mis-joins were corrected manually according to Hi-C contact map and global alignment resulted from HaploMerger2 (hm.new_scaffolds). Hicexplorer v3.4.1 (Wolff et al. 2018) was used to calculate Hi-C links of read pairs to each haplotype. To guide scaffolds to chromosomes, telomeres were identified by telomere_identification.pl (https://github.com/jimie0311/Puccinia-polysora-genome/blob/main/Figure-1/telomere_identification.pl). The possible tandem repeats, (CCTAAA/TTAGGG)n in most filamentous ascomycete fungi (Lue 2021), and other irregular type reported in closely related rust fungi, (CCCTAA/TTTAGG)n, (CCCCTAA/TTAGGGG)n, (CCCTAAA/TTTAGGG)n, (CCCTAA/TTAGGG)n, were all tested (Li et al. 2019; Tobias et al. 2021). Finally (CCCTAAA/TTTAGGG)n was identified as the telomere repeat units for *P. polysora*. The corrected scaffolds were rebuilt to chromosome level according to telomeres. The centromeres were identified by the cross-link signal based on Hi-C contact map (arrows in Figure 2B). The completeness of each haplotype was evaluated using BUSCOs of basidiomycota_odb9 and *Ustilago maydis* as the selected species for AUGUSTUS gene prediction (Stanke and Morgenstern 2005) in BUSCO v3.0.2 (Simão et al. 2015). The workflow of chromosome assembly has been illustrated in Figure 1A and the codes and pipelines used for the assembly pipeline were available in github (https://github.com/jimie0311/Puccinia-polysora-genome/blob/main/Figure-1/Figure-1A.md).

**FIGURE 1.**
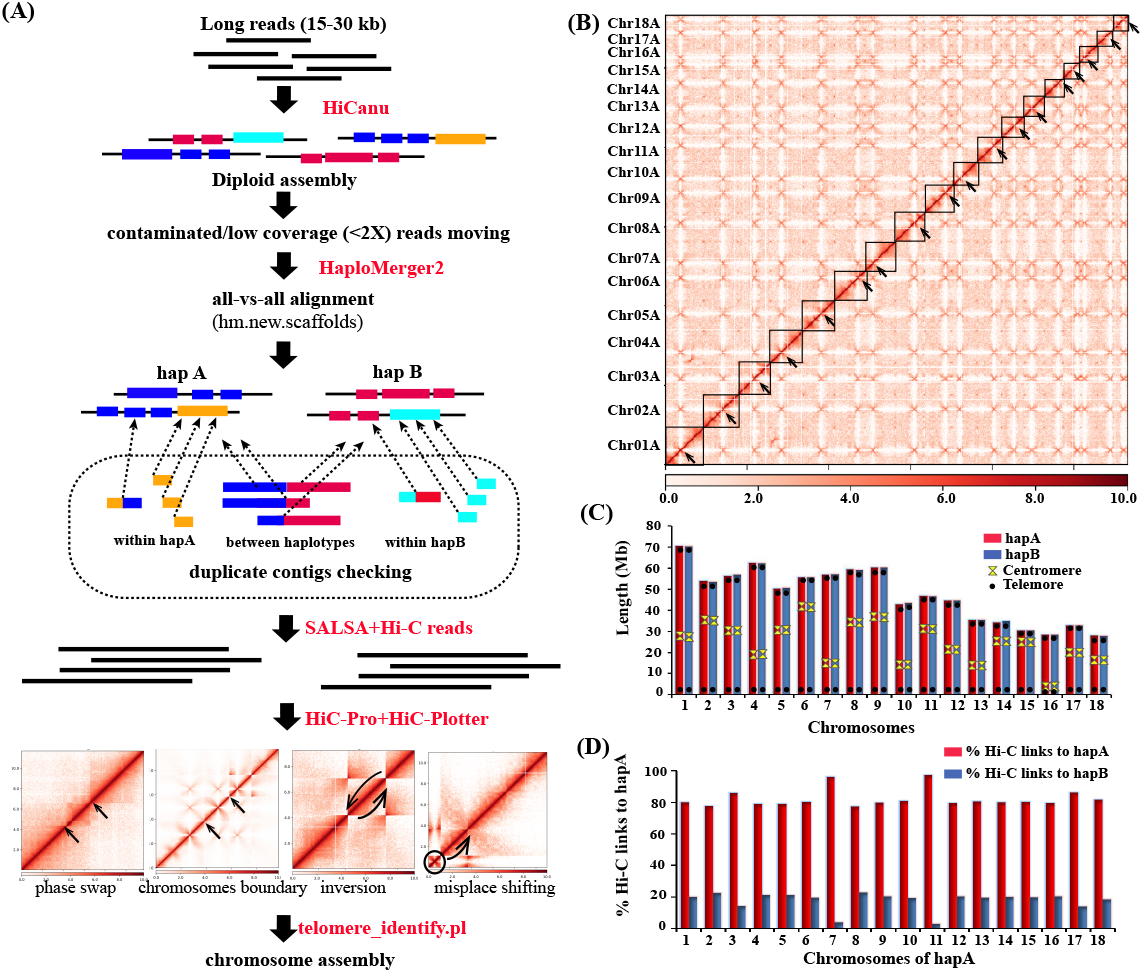
Chromosome-level assembly of GD1913 isolate of *Puccinia polysora*. (A) The dikaryotic phasing pipeline and switch strategies for duplicated contigs. Duplicated contigs were checked based on the all-vs-all sequence alignment from HaploMerger 2 (Huang et al. 2017). (B) Hi-C-based contig anchoring. The heat map showed the density of Hi-C interactions within haplotype A. The 18 chromosomes are highlighted by blue squares with arrows pointed at centromeres. (C) Schematic representation of assembled chromosomes for GD1913 of each haplotype. (D) Percentage of Hi-C links of each chromosome of haplotype A to either haplotype A (blue) or haplotype B (red).

**FIGURE 2.**
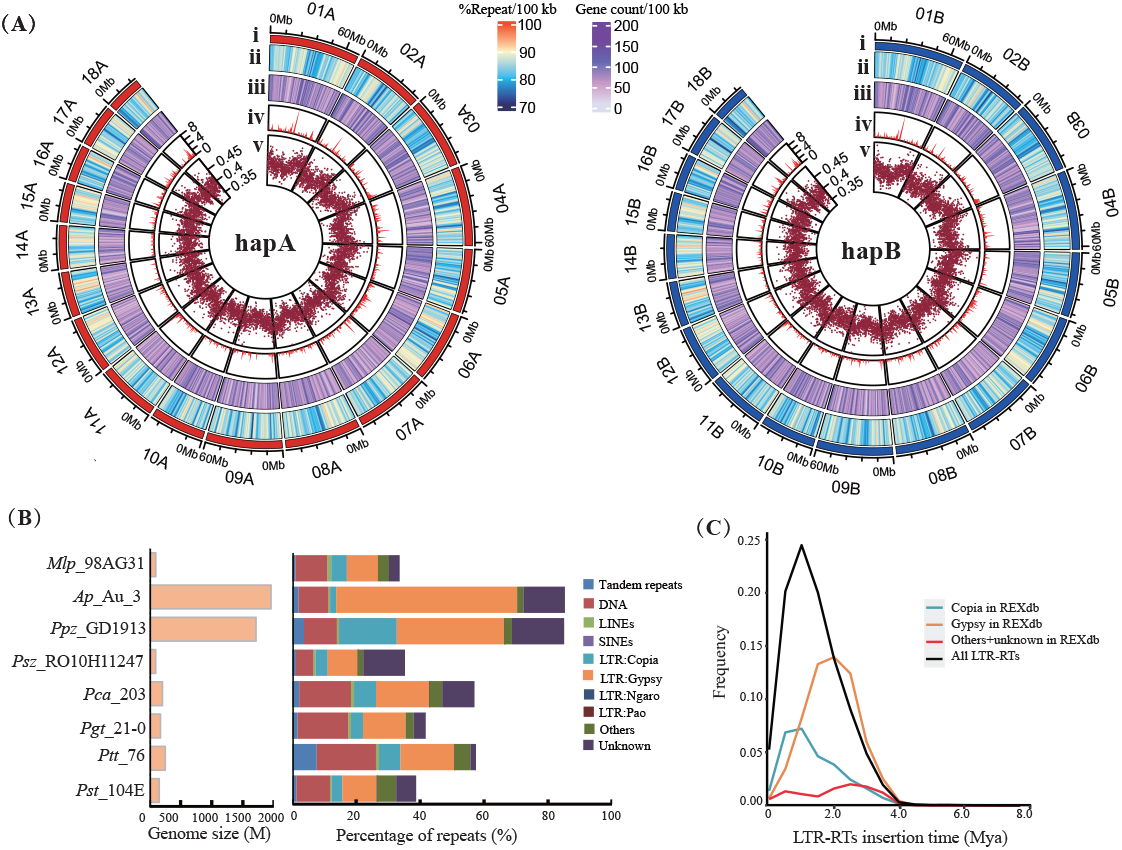
Genomic features of *P. polysora* f.sp. *zeae* isolate, GD1913 and repeat comparison analyses with its close relatives. (A) Genomic landscape of 18 chromosomes for haplotype A (hapA) and haplotype B (hapB). From outer to inner circles: (i) Chromosomes, (ii) Repeat density, (iii) Gene density, (iv) Candidate effector density, (v) Guanine-cytosine (GC) content. The tracks from ii to iv are estimated by non-overlapping 100 kbp sliding windows. (B) Comparison of genome size (two nuclei) and repeat contents among *Puccinia polysora* f.sp. *zeae* (*Ppz_*GD1913) and other rust species, *Melampsora larici-populina (Mlp*_98AG31), *Austropuccinia psidii (Ap_Au_*3), *Puccinia sorghi* f.sp. *zeae (Psz__*RO10H11247), *Pucccnia coronata* f.sp. *avenae (Pca_203), Puccinia graminis* f.sp. *tritici (Pgt_21-0), Puccinia triticina* f.sp. *tritici (Ptt*_76) and *Puccinia striiformis* f.sp. *tritici* (*Pst_*104E). (C) The insertion time (Mya) distribution of intact LTR-RTs in *P. polysora*.

### 2.4 RNA extraction and sequencing

For gene annotation, transcriptome sequencing of *P. polysora* from various infection time points was performed. Corn seedlings of 18 pots (10 seedlings/pot) were inoculated with 10 mg spores/mL mixed of Tween 20 (v/v: 0.05%). About five infected leaves were cut off per biological replicate at 1 day post inoculation (dpi), 2 dpi, 4 dpi, 7dpi, 10 dpi, and 14 dpi. In addition, germinated spore samples were prepared by placing 30 mg of fresh urediniospores on the surface of sterile water at 25–28°C in dark for12 hours. Three biological replicates were performed for each condition. All samples were collected in 4 mL Eppendorf tubes, frozen in liquid nitrogen before storing at −80°C. Before RNA extraction, samples were ground in liquid nitrogen and RNA was extracted using the RNeasy Plant Minikit (Qiagen) according to the manufacturer’s protocols (https://www.qiagen.com/us/resources/download.aspx?id=246847e7-0095-43e4-8d1d-41df3f9153dd&lang=en). After checking the RNA quality by Bioanalyzer, ~350 bp library was constructed and sequenced by Illumina NovaSeq 6000 platform (Annoroad Gene Technology Co., Ltd, Beijing, China). Before transcriptome assembly, raw reads were trimmed using Trimmomatic v0.36 (Bolger et al. 2014) with the settings: ILLUMINICLIP 2:30:10 LEADING 3, TRAILING 3 SLIDINGWINDOW 4:10 and MINLEN 50. A reference-guided transcriptome assembly was performed in Trinity v2.8.5 (Grabherr et al. 2011) using combined reads from germinated spores and infected leaves.

### 2.5 Repeats annotation and LTR aging

To compare the repeat contents between *P. polysora* (*Ppz*_GD1913) and other rust species, the genome data of *Melampsora larici_populina (Mlp_98AG31), P. sorghi (Psz_*_RO10H11247), *P. coronata* (Pca203), *P. graminis* (*Pgt*21-0), *P. triticina (Pt76)* and *P. striiformis* (*Pst*104E) were downloaded from NCBI or MycoCosm (DOE Joint Genome Institute), and *Austropuccina psidii* (*Ap*_Au_3) was downloaded from European Nucleotide Archive (ENA) at EMBL-EBI. The repeat contents were identified by a merged library including a self library in RepeatMasker v4.0.8 (Smit et al. 2015) and a genome-trained library from RepeatModeler v1.0.11 (Smit and Hubley 2008). The retrotransposons with long terminal repeats (LTR-RTs) of GD1913 were identified with an in-house pipeline that uses *LTRharvest* (Ellinghaus et al. 2008) packaged in LTR_retriever v2.9.0 (Ou and Jiang 2018). The classification of LTR-RTs was annotated based on REXdb (http://repeatexplorer.org) (Neumann et al. 2019) performed by TEsorter (Zhang et al. 2022).The genetic distances (GD) of each pair of LTR-RTs were calculated by *dist.dna* in R package of *ape*. The insert times (T) of LTR-RTs were estimated following the formula, T = GD/(2*mutation_rate). We used a mutation rate of 2.0×10^-8^ from a closely related fungus, *Schizophyllum commune* (Baranova et al. 2015).

### 2.6 Gene prediction and genome annotation

The two haplotypes were annotated independently using Funannotate v1.8.7 (https://github.com/nextgenusfs/funannotate/releases/tag/v1.8.7). A pipeline of core modules was applied as mask-> train->predict->update-> fix->annotate. The repeats of the assembled genome were soft-masked according to the results of 2.5. The prediction step (funannotate predict) was run with transcript evidence from Minimap 2 (Li 2018) RNA-seq alignments and genome-guided Trinity assemblies. Transcript evidence was aligned to two haplotypes separately and the protein evidence was aligned to the genome via Diamond (Buchfink et al. 2015)/Exonerate (Slater and Birney 2005) with the default UniProtKb/SwissProt protein database (http://legacy.uniprot.org/uniprot/?query=reviewed%3Ayes) from funannotate. The PASA gene models were parsed to train AUGUSTUS v3.2.3 (Stanke and Morgenstern 2005), snap and GlimmerHMM. In addition, GeneMark_ES (Lomsadze et al. 2005) was self-trained using two haplotypes’ sequences. All above evidence was combined with default weight settings using Evidence Modeler (Haas et al. 2008) and filtered by removing genes with short length (< 50 aa), spanning gaps and transposable elements. The tRNA genes were predicted using tRNAscan-SE v1.3.1 (Lowe and Chan 2016). Funannotate update command to add UTR data to the predictions and fix gene models that are in disagreement with the RNA-seq data. Functional annotation was performed based on available databases including Pfam v34.0) (Finn et al. 2014), InterPro (v86.0) (Jones et al. 2014), eggNOG (v5.0) (Huerta-Cepas et al. 2019), UniProtKB (v.2021_03), MEROPS (v12.3) (Rawlings et al. 2016), carbohydrate hydrolyzing enzymatic domains (CAZymes) (Terrapon et al. 2017) and a set of transcription factors based on InterProScan domains to assign functional annotations.

### 2.7 Interhaplotype variation analysis

Small variants including SNPs and indels were identified by mapping trimmed short reads from DNA against haplotype A with BWA-MEM v0.7.8 (Li and Durbin 2010). After removing PCR duplicates using Picard v2.18.27 (https://github.com/broadinstitute/picard), the bam file was input to GATK 4.1.9 (https://github.com/broadinstitute/gatk) to call SNPs. SNPs and Indels were filtered using gatk VariantFiltration with parameters QD > 2.0 & MQ > 40.0 & FS < 50.0 & SQR <3.0 for SNPs and QD > 2.0 & QUAL > 30.0 & FS < 200.0 & ReadPosRankSum>-20.0. Variants were annotated from genome location and functional impact using SnpEff v4.3 (Cingolani et al. 2012). To estimate structure variations (SVs) between two haplotigs, we aligned 18 chromosomes of haplotype A to their corresponding chromosomes in haplotype B by using the nucmer program in Mummer 4 (-maxmatch -t 100 -l 100 -c 500) (Marçais et al. 2018). The delta file was filtered by parameters, delta-filter -i 95 -l 1000. To make sure the filter parameters acceptable, *show-coords* and *show-aligns* were used to check alignment details. Finally, the chromosome-pair synteny was visualized by *mummerplot*. In addition, the chromosome-pair alignments were also analyzed using Assemblytics (http://assemblytics.com/) to estimate SVs of three major categories, including insertion/deletion, tandem repeats contraction/expansion and repeats contraction/expansion. The running parameters of Assemblytics were set with the unique sequence length=10,000, maximum variant size=10,000 and minimum variant size=1. To identify the orthologous genes between two haplotypes, we used OrthoFinder v2.5.4 with the default parameters (Emms and Kelly 2015).

### 2.8 Prediction of candidate effectors

Secreted proteins were predicted based on two rules 1) the presence of a predicted signal peptide using SignalP 4.0 (Petersen et al. 2011); and 2) the absence of predicted transmembrane domains outside the first 60 amino acids (with TMHMM 2.0). To predict candidate effectors, we use EffectorP 3.0 (Sperschneider and Dodds 2021), which applied two machine learning models trained on apoplastic and cytoplasmic effectors. The density plots for genes, repeats, and secreted proteins from chromosomes in two haplotypes were generated using KaryoploteR (Gel and Serra 2017) with the scripts available at https://github.com/jimie0311/Puccinia-polysora-genome/blob/main/Figure-4/Figure-4B.md. A recent study identified the first effector of *P. polysora*, *AvrRppC* (Deng et al. 2022). To locate its genome position, we blasted the CDS sequence of *AvrRppC*^ref^ against genome sequences of two haplotypes in this study.

To understand the expression pattern of secreted proteins, we used all expression data from germinated spores and six time-points to perform a differential expression analysis. The allele specific mapping of Kallisto v0.48 (Bray et al. 2016) was applied to obtain the expression levels in transcripts per million (TPM) of genes on both haplotypes. FeatureCounts v1.5.3 (Liao et al. 2014) was used to generate read counts for each gene model of secreted protein. Differentially expressed genes were identified by expression in plants relative to germinated spores (|log fold change| > 1.5; adjusted *P* < 0.1) using the DESeq2 R package (Love et al. 2014). The average rlog-transformed values for each gene were used for clustering using the *k*-means method. The optimal number of clusters was defined using the elbow bend method (https://www.delftstack.com/howto/r/k-means-clustering-r) and circular heatmaps were plotted using the Circlize R package (Gu et al. 2014).

### 2.9 Comparative genome analysis

The high quality of *P. polysora* genome provides the opportunity for comparison with other chromosome-level references, e.g., *Pgt*21-0 (Li et al. 2019), *Pt*76 (Duan et al. 2022) and *Pca203* (Henningsen et al. 2022). Syntenic gene pairs among *Puccinia* species were identified using the MCSCAN toolkit (Wang et al. 2012). Figures were plotted using MCscan (Python version). (https://github.com/tanghaibao/jcvi/wiki/MCscan-(Python-version). Orthofinder v2.5.4 (Emms and Kelly 2019) was applied to identify the orthogroups of six rust species with available haplotype genomes. Besides the above three *Puccinia* species and *P. polysora* in this study, the primary haplotype of *P. striiformis* (*Pst*104E) and *Austropuccinia psidii* (Au_3) were also included. The phylogenetic relationship of these species was visualized by FigTree v1.4.3 (http://tree.bio.ed.ac.uk/software/figtree/). The gained (+)/lost (−) gene numbers were calculated by comparing the two most close species according to Orthogroups.GeneCount.tsv in Orthofinder output.

### 2.10 Population genetic analyses

To understand the population differentiation of *P. polysora* in China, 79 isolates were resequenced. Reads were qualified by Trimmomatic v0.36 (Bolger et al. 2014) using the parameters described in 2.4 RNA extraction and sequencing. To assess whether each isolate comprises a single genotype free of contamination, the distribution of read counts for bi-allelic SNPs was calculated by R package vcfR v1.8.0 (Knaus and Grunwald 2017) and plotted using ggplot2 (v2.2.1) (Wickham 2016). The normal distribution of read allele frequencies at heterozygous positions is expected to rule out contamination from other *Ppz* genotypes and only the pure isolates were applied in the following population genomic analyses. All clean reads of pure isolates were mapped to haplotype A and SNPs calling and filtration were performed by GATK v4.1.9 as described in 2.7 Interhaplotype variation analysis section. Population structure was detected and quantified using principal component analysis (PCA), which was performed using Plink v1.9 (Purcell et al. 2007). Pairwise FST values were estimated between geographical regions using the PopGenome R package (v2.6.1) (Pfeifer et al. 2014). To assess the effects of variants, the vcf file including selected SNPs was run in SnpEff v4.3 (Cingolani et al. 2012). Because the sexual stage of *P. polysora* has not been reported, we used the standardized index of association (*rd*) to test linkage disequilibrium in the Chinese *Ppz* population. We constructed 100 sets of 10,000 random SNPs by samp.ia function from R package *poppr* (Kamvar et al. 2014) to generate a distribution of *rd* values. The observed *rd* distribution of the Chinese population was compared to the distribution of 10,000 *rd* values constructed using fully randomly simulated datasets with 0%, 50%, 75% and 100% linkage.

## 3 RESULTS

### 3.1 Chromosome-level genome assembly and haplotype-phasing of *P. polysora*

We generated a total of 53 Gbp of circular consensus reads (32× coverage) from single-molecule real-time sequences on the PacBio Sequel II. A total of 4056 contigs were assembled with a genome size of 1.76 Gbp and a contig N50 of 1.9 Mbp (Table 1). A total of 1627 contigs, accounting for ~2% of the *de novo* assembly size were excluded due to small size, low coverage or high mitochondrial similarity (Table S2). By using Haplomerger2, the remaining contigs were assigned to two haplotypes, A and B, with 613 contigs representing ~862 Mbp and 1321 contigs representing ~852 Mbp, respectively. These contigs were further connected to 173 and 344 scaffolds in haplotypes A and B by Hi-C scaffolding (Table 1), after manual checking of 33 duplicated scaffolds between (17) and within (16) haplotypes (Table S3). By considering both Hi-C contact maps and the all-vs-all alignment from Haplomerger2, 18 super scaffolds were binned for each haplotype manually. The Hi-C contact map showed evidence of a single centromere on each scaffold and all scaffolds contained telomere sequences at either end (Table 1, Figures 1B, 1C), indicating that each haplotype contains 18 chromosomes. This is consistent with the observations of 18 chromosomes in the genome of other *Puccinia* species (Boehm et al. 1992; Li et al. 2019; Wu et al. 2021; Duan et al. 2022). In addition, 15 (6.5 Mbp) and 20 (4.5 Mbp) scaffolds in haplotype A and B respectively could not be correctly connected to any chromosomes due to short length or insufficient Hi-C signal. The 18 chromosomes of haplotype A showed an average of over 80% Hi-C links to haplotype A, suggesting high levels of nuclear phasing (Figure 1D). The chromosomes were numbered according to the synteny with the other *Puccinia* species (see below). In addition, a cyclized mitochondrial sequence (67 Kb) was assembled.

**Table 1.** Assembly information and completeness evaluation of *Puccinia polysora* f.sp. *tritici*, GD1913.

| Parameters | before haplotype-phased |  |
| --- | --- | --- |
| No. of contigs | 4056 |  |
| No. of bases (Gbp) | 1.76 |  |
| Size of longest contig (Mbp) | 14.7 |  |
| N50 (Mbp) of contigs | 1.9 |  |
|  | after Hi-C scaffolding |  |
|  | Haplotype A | Haplotype B |
| No. of scaffolds | 173 | 344 |
| Size of scaffolds (Mbp) | 862 | 855 |
| Size of longest scaffold (Mbp) | 31 | 39 |
| N50 (Mbp) of scaffolds | 13.2 | 9.5 |
|  | after Hi-C contact map binning |  |
| Total size (Mbp) | 856 | 855 |
| No. of chromosomes | 18 | 18 |
| No. of unplaced scaffolds | 15 | 20 |
| Size of chromosomes (Mbp) | 849 | 850 |
| Size of longest chromosome (Mbp) | 70.5 | 70.3 |
| GC content | 40.1% | 40.0% |
| Complete BUSCOs | 89.7% | 89.7% |
| Complete and Single-copy BUSCOs | 88.6% | 88.6% |
| Complete and Duplicated BUSCOs | 1.1% | 1.1% |
| Fragmented BUSCOs | 3.5% | 3.5% |
| Missing BUSCOs | 6.8% | 6.8% |

### 3.2 Repeats in the supersize genome of *P. polysora*

The completeness of the *Ppz* assembly was assessed based on highly conserved Basidiomycete genes (Badidiomycete_odb9, 1335 BUSCOs), which suggested that the two haplotypes showed a similar level of completeness, with approximately 90% complete BUSCO genes and an additional 3.5% fragmented BUSCOs, respectively (Table 1). However, 126 BUSCO genes were complete in haplotype A but not present in haplotype B. On the contrary, 127 complete BUSCO genes are only present in haplotype B, which suggests interhaplotype variation in *P. polysora*. Considering the two haplotypes in combination, only 11 BUSCOs were missing or fragmented, resulting in 99% complete BUSCOs. That means we assembled a near-complete genome of *P. polysora*.

Using the *de novo* library of RepeatMasker and genome-trained library of RepeatModeler, RepeatMasker detected repeats accounting for 85% of the diploid assembly (both haplotypes), which is significantly higher than observed in closely related species of *Puccinia* (30%–59%) (Miller et al. 2018; Schwessinger et al. 2018; Wu et al. 2021), but similar to *Austropuccinia psidii* (85% estimated in this study but 91% in its original description) (Tobias et al. 2021). The repeat density and low GC content (~40%) along the two haplotypes were illustrated in Figure 2A. The large percentage of repeats in *P. polysora* was mainly caused by Class I retrotransposons with long terminal repeats (LTR), with the LTR-Gypsy superfamily most abundant (33.5%), and LTR-Copia next (17.9%). The TE family composition was similar to other rust species, except for *A. psidii*, which is LTR-Gypsy dominated with LTR-Copia accounting for only 1.9% (Figure 2B). In addition, about 20% and 16% of repeats of *P. polysora* in two haplotypes were unclassified repeat families based on Repbase (Bao et al. 2015).

Since LTR-RTs occupied the majority of the *P. polysora* genome, we examined whether these repetitive sequences slowly accumulated over time or alternatively were subject to sudden expansion in the life history of *P. polysora*. A total of 41,634 LTR-RT pairs were extracted. An LTR-RT burst was estimated at around 1.7 Mya (Figure 2C).

### 3.3 High levels of interhaplotype gene content and structural variation

RNA-seq reads from germinated spores and infected corn leaves at 1, 2, 4, 7, 10, and 14 dpi (Table S4) were pooled and used to generate genome-guided transcriptome assemblies. In total, we annotated 23,270 and 24,242 gene models on haplotype A and haplotype B, respectively (Table 2). The protein and CDS sequences are available from https://github.com/jimie0311/Puccinia-polysora-genome. The gene space of these genes accounts for only ~4% of the total genome size. The two haplotypes showed a similar level of functional annotation with about 51% of proteins having at least one functional annotation. Orthofinder identified a total of 13,057 common orthogroups between the two haplotypes, involving 20,802 genes from haplotype A and 21,519 genes from haplotype B, over 90% in each haplotype. In addition, 198 and 235 orthogroups were specific to haplotype A or B respectively. About 70% of haplotype-specific genes have no functional annotations, with the remainder having annotated functions mainly in RNA mediated transposition, transmembrane amino acid transposition or ATP binding (Table S5).

**Table 2.**
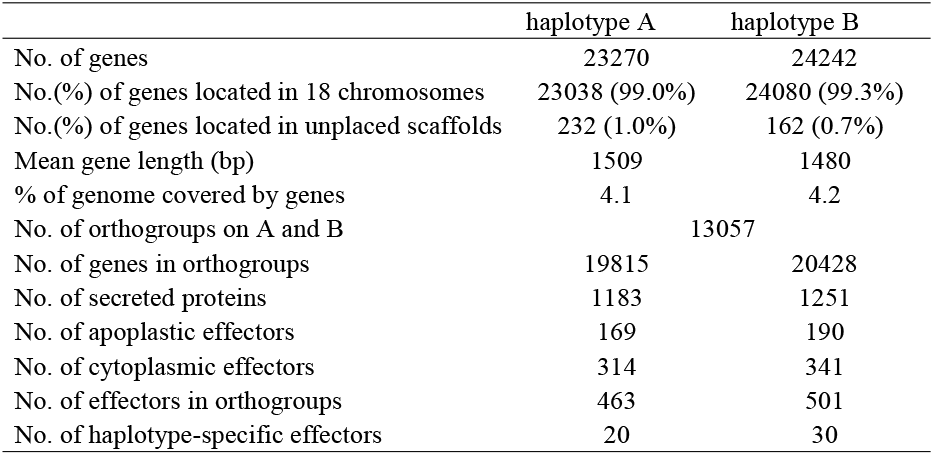
Genes, orthogroups, and effectors for two haplotypes of *Pucciniapolysora* f.sp. *tritici*, GD1913.

By mapping Illumina reads to only haplotype A, we detected a total heterozygous rate of 6.1 variants/kbp, in which the SNP variant rate is dominated (5.8 SNPs/kbp). About 78% of the SNPs were located in intergenic regions and the rest (22%) were detected in 22,802 proteincoding genes. The results of Assemblytics suggested that structural variation between haplotype A and B comprised about 3.0% (26/849 Mbp) of the haploid genome size (Figure 3A). The full chromosome-pair alignment between two haplotypes is illustrated in Figure 3B, with alignments of chromsome09 and chromsome15 in Figure 3C and D, showing some insertions/deletions and inversions, respectively. Among three types of variation, insertions/deletions and repeat expansions/contractions are more prevalent than tandem expansions/contractions. Besides, variation with size bins of 500 to 10,000 bp is the most prevalent, which accounts for 2.8%. Because the maximal variant size was restricted to 10,000 in Assemblytics, the actual difference between the two haplotypes could be even higher than estimated.

**FIGURE 3.**
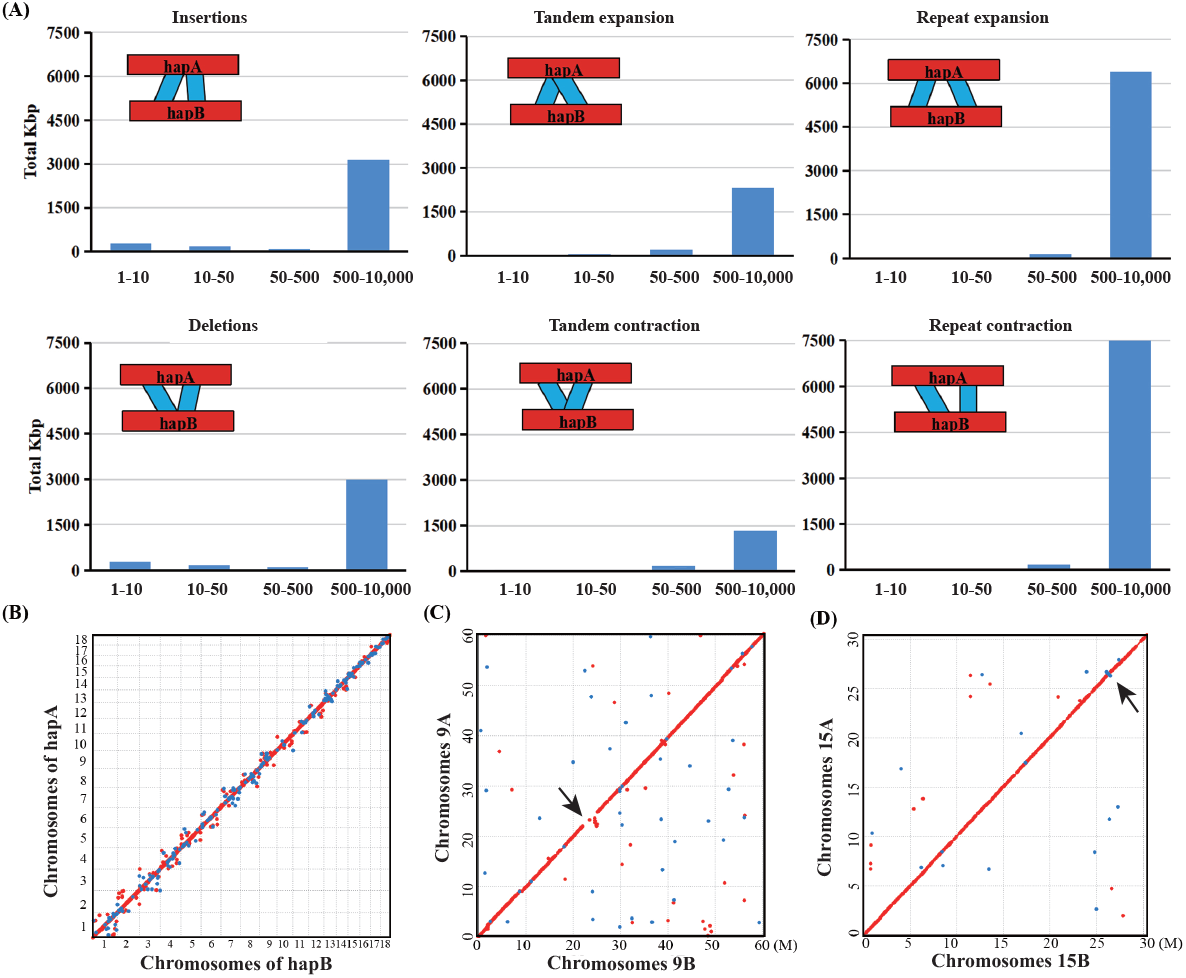
The interhaplotype variation of *Ppz*-GD1913 isolate. (A) Summary of interhaplotype variation between haplotype A and B using Assemblytics. Six types of specific variation were illustrated by schematic plot and each bar chart indicates the number of bases categorized to each type. (B) Synteny plot of haplotype A and B using Mummer. (C) and (D) Full-chromosome alignments of chromosomes 9 and 15 showing insertion/deletion and inversion (arrows) between haplotypes.

### 3.4 Conservation of expression patterns between secreted proteins from two haplotypes

We predicted 1183 and 1251 secreted proteins on haplotype A and haplotype B, respectively. By using the machine-learning tool, EffectorP v3.0, about 14% and 27% of all secreted proteins were predicted as apoplastic and cytoplasmic effectors, respectively (Table 2). The protein sequences of secreted proteins are available from https://github.com/jimie0311/Puccinia-polysora-genome. GO enrichment analysis and Interproscan annotation predicted some effectors involved in hydrolase activity, catalytic activity, protein tyrosine phosphatase activity and metal ion binding, but most effectors had no homology with known or predicted functions (Table S6, Figure S2). Then we used gene expression data to predict clusters in the secretome that are differentially expressed during infection. The elbow bend plot (Figure S4) suggested the total within-group sum of squares tend to be level off. Therefore, the expression data of secreted proteins were assigned to seven clusters. Genes in cluster 1 showed high expression in germinated urediniospores and early infection (1dpi, 2dpi) but low expression at 4dpi and 7dpi when haustoria form. On the contrary, genes in clusters 2, 3, 4 and 5 showed low expression in germinated spores, and highest in planta expression at days 1-2 (cluster 2), days 2-7 (cluster 3) and from days 4-7 (cluster 4) and days 7-14 (cluster 5) Clusters 6 and 7 were more uniform in expression through these stages, although cluster 6 genes increased later in infection. About 24% to 67% of the secreted proteins in these clusters were predicted as candidate effectors by EffectorP (Table S7). A similar set of expression profile clusters were also detected for genes in haplotype B (Figure 4A). Nevertheless, two haplotypes presented different levels of carbohydrate-active enzymes (CAZymes). A total of 209 and 298 CAZymes were detected, of which 9 vs 79 CAZymes were predicted to be secreted in haplotypes A and B, respectively (Table S7). Among CAZymes subclasses, Glycoside hydrolase (GH) enzymes are abundant, accounting for 48% (88 GH families) and 42% (126 GH families) of all CAZymes of haplotype A and haplotype B, respectively. Of these, 51 GH families were predicted to be secreted. The GH5 (cellulase and other diverse forms being exo-/endo-glucanases and endomannannases) family (Langsto et al. 2011) was observed to be largely expanded in *P. polysora* as well as other *Puccinia* species (Figure 5A). Besides, AA3 (glucose-methanol-choline oxidoreductases), CE4 (chitin and peptidoglycan deacetylases), GH18 (chitinases of classes III and V), CH47(α-mannosidases), GT2 and GT90 are also abundant in *P. polysora*.

**FIGURE 4.**
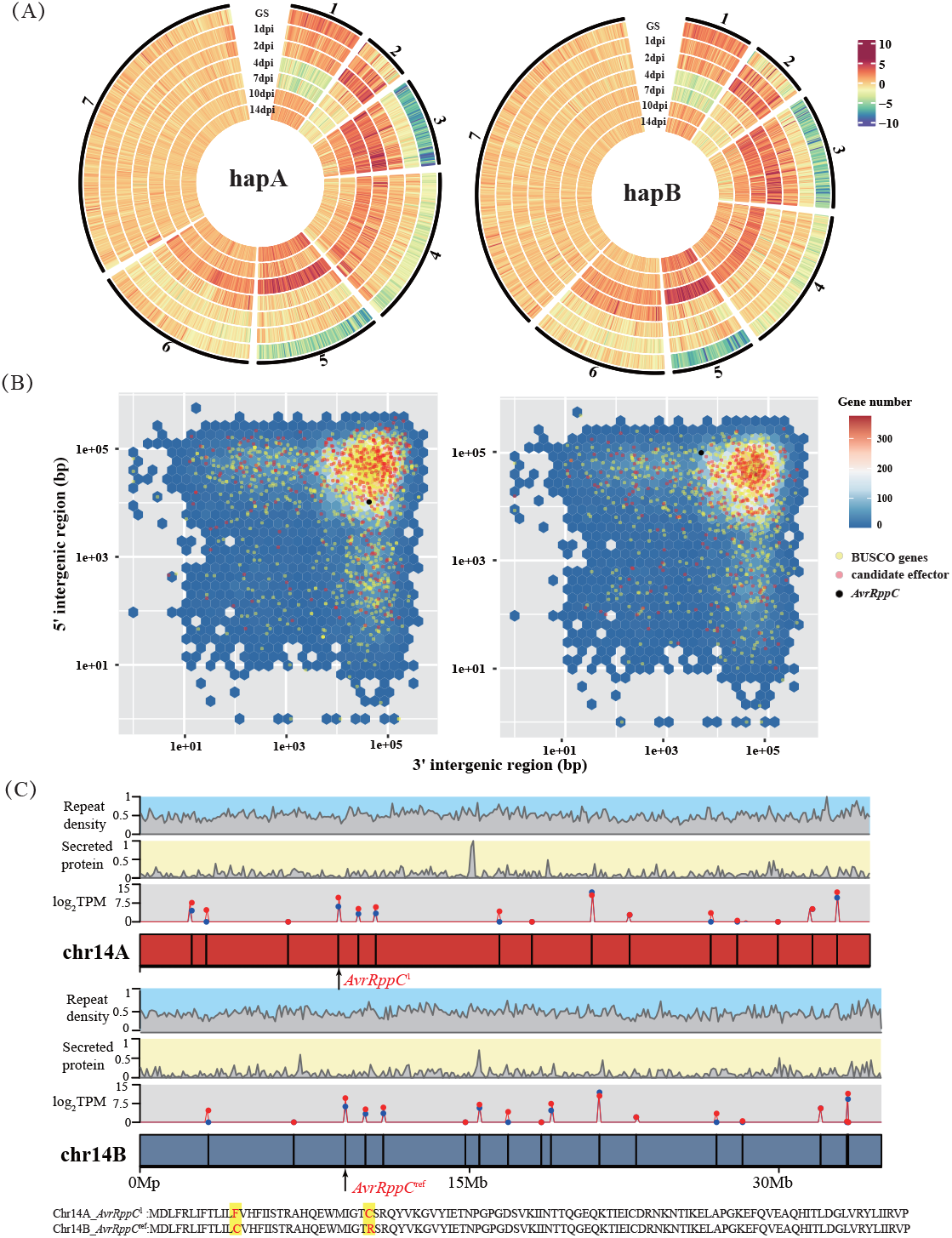
Analyses of gene expression and gene distance of predicted secretome of *P. polysora*. (A) Clustering analysis of gene expression of secretome on two haplotypes. Heatmaps show rlo- transformed expression values. Cluster numbers are shown outside the graphs, and tracks represent gene expression in germinated spores (GS), and infected tissues at 1 dpi, 2 dpi, 4 dpi, 7 dpi, 10 dpi and 14 dpi. (B) Hexplots for closest-neighbor gene distance density of *P. polysora* in haplotype A (left) and haplotype B (right). Circle dots represent two gene categories, BUSCOs (yellow) and predicted candidate effectors (red). The black dots represent the distribution of the avirulent gene, *AvrRppC*. (C) Gene and repeat density as well as log2TPM (transcripts per million) of candidate effectors from chromosomes 14A and 14B. The red and blue points represent expression values of candidate effectors at 1 dpi and 7dpi. Positions and protein sequences of *AvrRppC* genes are presented.

**FIGURE 5.**
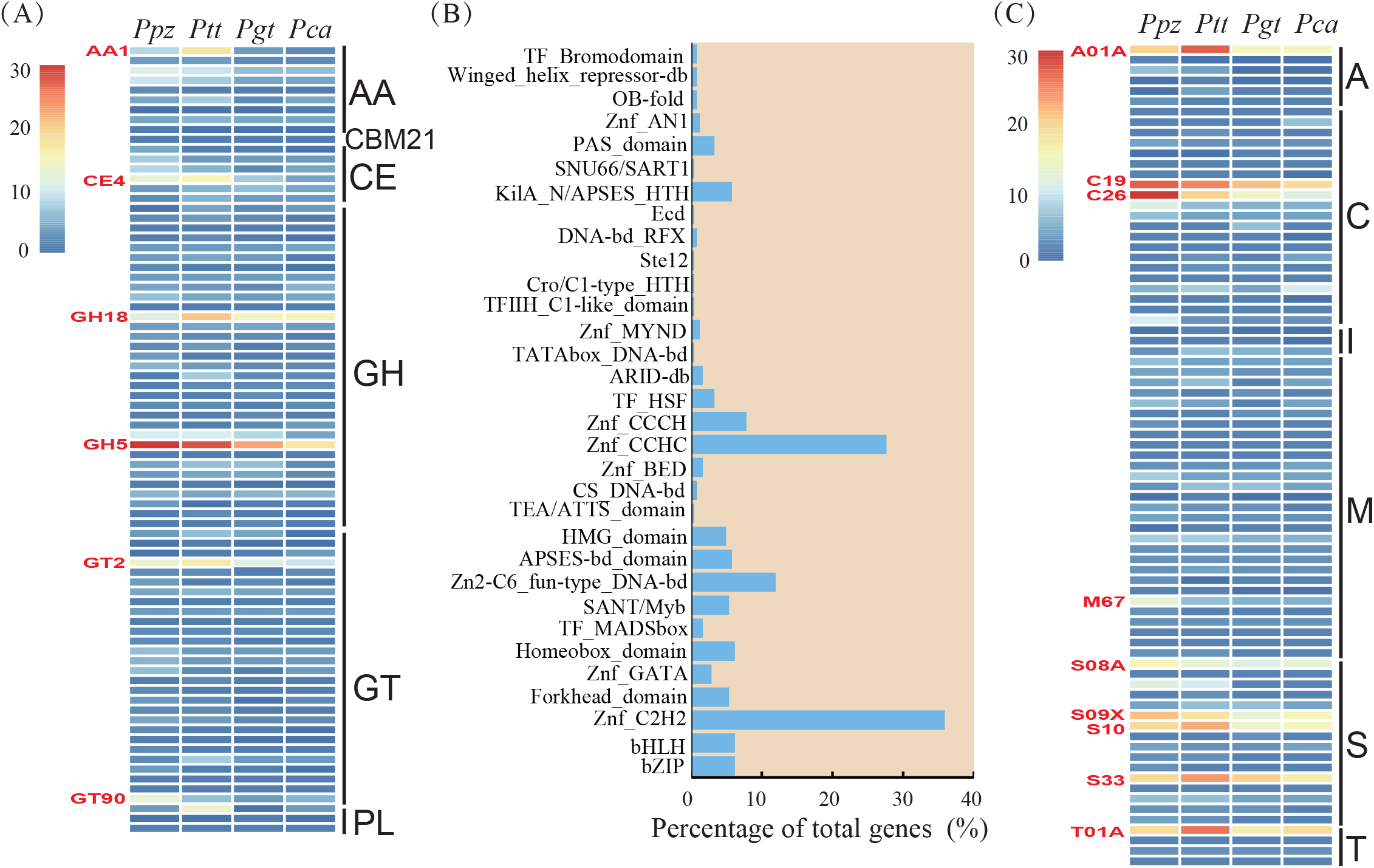
Functional annotation of CAZymes, transcription factors and Merops proteases of *Puccinia polysora* GD1913 (*Ppz*-GD1913) and its comparison with other *Puccinia* species. (A) CAZyme families comparison of *P. polysora (Ppz*, GD1913), *P. triticina (Ptt, Pt76), P. graminis (Pgt, Pgt*21-0) and *P. coronata (Pca, Pca203)*. Heat maps showing gene numbers annotated in the following classes: auxiliary activities (AA), carbohydrate-binding modules (CBM), carbohydrate esterases (CE), glycoside hydrolases (GH), glycosyltransferases (GTs), and polysaccharide lyases (PL). Expanded families of AA1, CE4, GH18, GH5, GT2 and GT90 were highlighted in red. (B) Percentages of genes from *P. polysora* are predicted to encode members of various fungal transcription factor classes based on InterProScan and Eggnog annotation. (C) Merops families comparison of *Ppz, Ptt, Pgt* and *Pca*. Heat map showing gene numbers annotated in the classes of aspartic acid (A), cysteine(C), peptidase inhibitors (I), metalloprotease (M), serine protease (S), and threonine protease (T) which were labeled in the right of the heat map and expanded families were highlighted in red.

We also investigated other factors potentially associated with infection or plant immune inhibition, such as transcription factors (TFs) and peptidases. We found most TF families of *P. polysora* have low abundance except C2H2-type (IPR013087) and CCHC-type (IRP001878) zinc finger class, which have 87 and 67 members in haplotypes A and B (Figure 4B). In total, we annotated 286 and 304 proteases in haplotypes A and B, with A01A (aspartic proteases), C19 (ubiquitinyl hydrolases), C26 (gamma-glutamyl hydrolase), S08A (subtilisin-like serine protease), S09X (glutamyl endopeptidase C), S10 (carboxypeptidase Y), S33 (prolyl aminopeptidase) and T01A (component peptidases of the proteasome) families expanded in *P. polysora* as well as in other three *Puccinia* species (Figure 5C). A total of 8 (2.8%) and 30 (10.0%) proteases were predicted to be secreted and these proteases showed a discrete distribution in each cluster without an obvious clustering pattern (Table S7).

### 3.5 Density of candidate effectors and *AvrRppC* anchoring

Analysis of the local gene density, measured as flanking distances between neighboring genes showed that the flanking distances in the *P. polysora* genome are generally rather far, with an average distance between genes of 100 kbp (Figure 4B). Accordingly, the surrounding genomic context of most genes in *P. polysora* genome is gene-sparse and repeat-rich. Large flanking distances are not specific to candidate effectors. In line with this pattern, the gene distance density plots revealed very similar distributions between all genes and candidate effectors, including the only known *Avr* effector gene, *AvrRppC* (Figure 4B). We further investigated whether candidate effectors present a different distribution of gene distance density compared to basidiomycete core ortholog genes of *P. polysora*. The results highlighted that candidate effectors are not located in peculiar gene-sparse areas, and the flanking distance centers of both BUSCOs and candidate effectors overlap with that of all genes (Figure 4B).

A previous study identified the avirulence effector, *AvrRppC*, and found six allelic variants which were named as *AvrRppC*^ref^, *AvrRppC*^A^, *AvrRppC*^C^, *AvrRppC*^E^, *AvrRppC*^F^, *AvrRppC*^J^, (Deng et al. 2022). By searching the CDS sequence of *AvrRppC*^ref^ against haplotypes A and B we anchored *AvrRppC* at ~ 9 Mbp on chromosome 14. Although the gene was not present in the original annotation, there was evidence of transcription from RNA-seq reads at 1 dpi and 7 dpi (Figure 4C). Additionally, we found the *AvrRppC* allele on Chromosome14B is *AvrRppC*^ref^ type whereas the allele in Chromosome 14A represents a new allele type, named *AvrRppC*^1^ which has two amino acids changes compared with *AvrRppC*^ref^ and is different from the other five previously reported variants.

### 3.6 Comparisons of synteny and orthogroups between *P. polysora* and its close relatives

Although *P. polysora* has a 7–10 times larger genome size than other rust species, the gene synteny amongst four *Puccinia* species with full chromosome assemblies is well conserved, with 18 chromosomes corresponding to each other clearly without any chromosome rearrangement (Figure 6A). A few large inversion blocks were detected in chromosomes 1, 3, 4, 7, 8, 9 and 10 between *PpzA* and *Pt*76 (Figures 6A, S3). Inversion blocks were also detected in chromosomes 5, 7, 10, 13 between *Pt*76 and *Pca*203, and chromosomes 2, 3, 5, 7, 8, 10 and 13 between *Pgt*21-0 and *Pca*203 (Figures 6A, S3).

**FIGURE 6.**
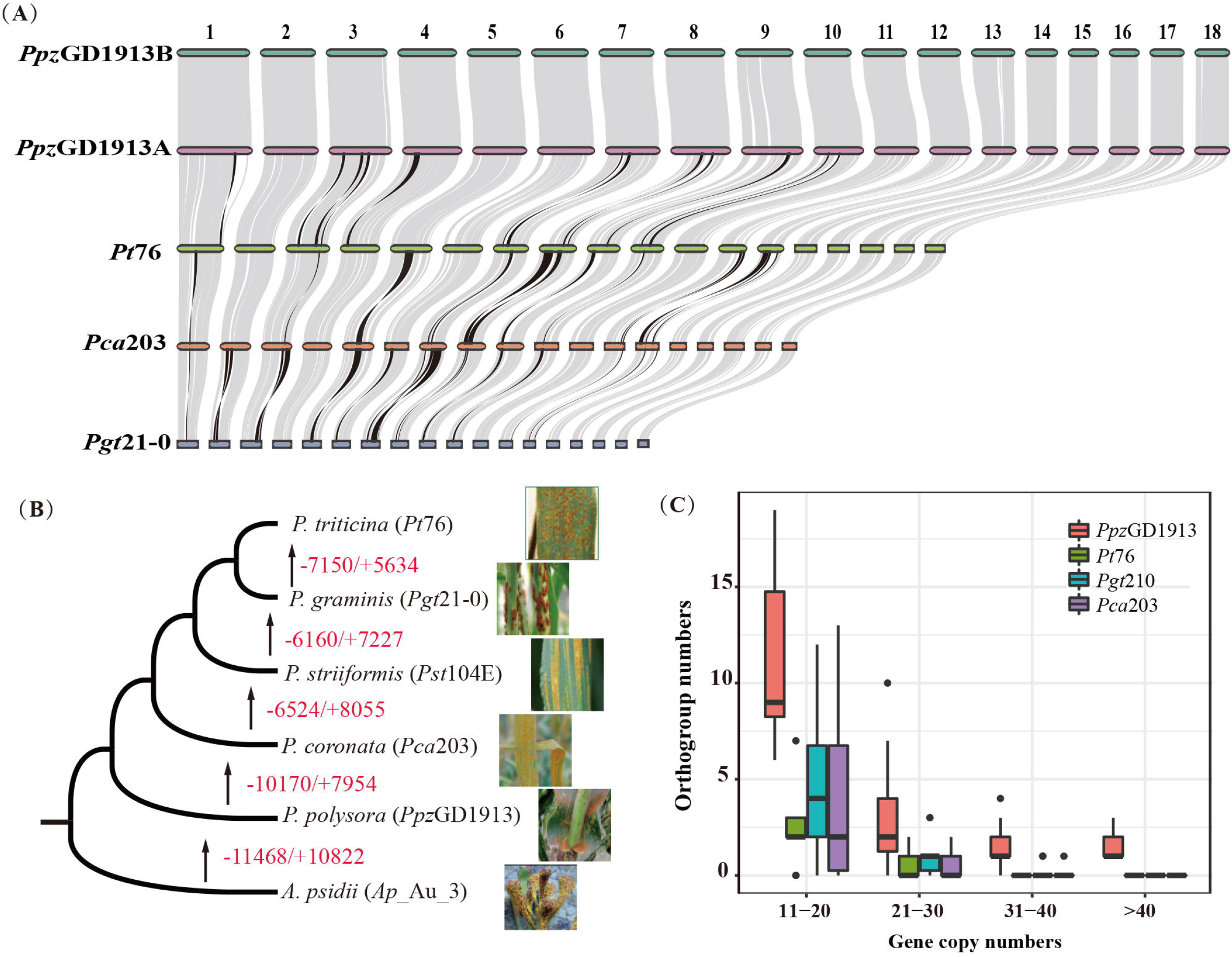
Comparative analyses of macro-synteny and orthogroups among *Puccinia. polysora* f. sp. *zeae* (*Ppz*GD1913A and *Ppz*GD1913B) and its close relatives, *Austropuccinia psidii* (Au_3), *P. triticina* f. sp. *tritici (Pt76), P. graminis* f.sp. *tritici* (*Pgt*21-0) and *P. coronata* f.sp *avenae* (*Pca*203). (A) Macro-synteny of four species in *Puccinia* with available haplotype- phased and chromosome-scale genome. Only the chromosomes from haplotype A/primary haplotype were compared. Grey lines are homologous proteins identified by MCscan Python version with default parameters and black lines present the genome inversions between species. For better visualization, the genome size ratio of *Ppz*B:*Ppz*A:*Pt*76:*Pca*203:*Pgt*21-0 = 0.2:0.2:1:1:1. (B) Phylogenetic relationship of *P. polysora* and other rust species was inferred by 1454 single-copy orthologue genes. The numbers indicated the sum of gained (+) and lost (−) genes from all orthogroups by comparing the two most close species. The typical symptoms caused by each species are listed accordingly. (C) Distribution of species-specific orthogroups with varied ranges of gene copy number among four *Puccinia* species.

The protein orthologs among four closely related *Puccinia* species were compared based on haplotype A of each species. The increased and reduced genes in all orthologs between two close species are labeled in the phylogenetic tree (Figure 6B). It was apparent that the reduced genes were much more than gained genes when compared from *P. polysora* to other *Puccinia* species. Therefore, *P. polysora* might experience gene expansion events compared with the other three species (Figure 6B, Table S8). Among the four species compared, a total of 11,840 orthogroups were identified. Genes of *P. polysora* were assigned to 7,148 orthogroups which was less than those of over 8,000 orthogroups for the other three *Puccinia* species. Nevertheless, *P. polysora* presented more species-specific orthogroups and these orthogroups are featured with a high copy number of gene duplication (Figure 6C). Based on Eggnog annotation result, the orthogroups of *P. polysora* with high copy numbers (>10) are mainly associated with the pathogenicity-related function, energy metabolism, RNA regulation and heat shock reaction (Table S9).

### 3.7 Population genomics analyses suggested lineage divergence and predominant asexual mode of reproduction

To understand the genetic structure of *P. polysora* in China, we analyzed genome resequencing data from 79 isolates of *Ppz* collected from across maize-growing regions of China. The analyses of bi-allelic frequency suggested the expected normal distribution of bi-allelic frequency for most isolates except GD1922-3 and GX1905-2 (Figure S5) which were excluded from the analysis. After removal, a total of 7,147,489 whole-genome SNPs were obtained from the remaining 77 isolates. PCA analysis indicated the presence of two genetic groups, with a major group consisting of 73 isolates (blue circle in Figure 7B) and a minor group containing only four isolates (red circle in Figure 7B). However, this separation was independent of the geographic origin of the isolates (North, Central, and South China). Although higher genetic differentiation was revealed between North China and South China populations (*Fst* = 8.1 ×10^-4^) than in other regional pairs (3.9×10^−4^ and 3.6×10^−4^), all pair-wise *Fst* values were close to zero indicating a lack of geographic differentiation. We used the standardized index of association, *rd*, to estimate the linkage disequilibrium of *P. polysora* population in China. The observed *rd* distribution for all isolates was between the values calculated from simulated datasets with 0% linkage, a sexual population, and 50% linkage (Figure 7C). The result suggested that, although *Ppz* population in China showed as a clonal population, its evolutionary history may be influenced by some extent of sexual reproduction.

**FIGURE 7.**
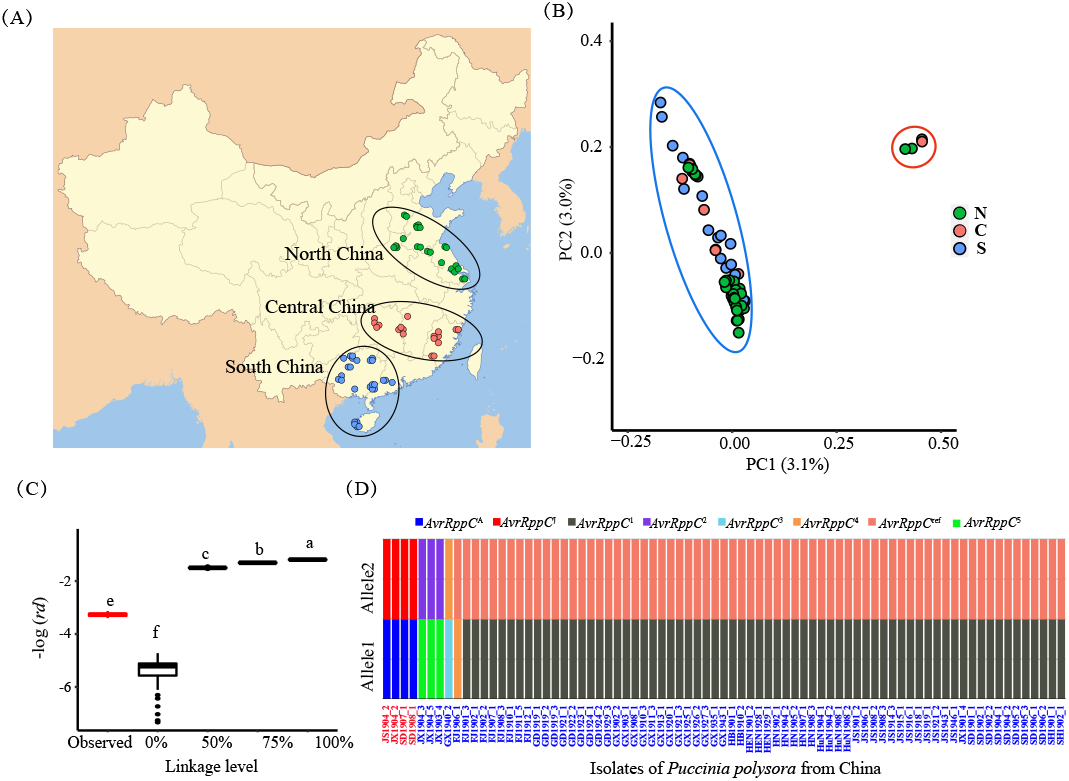
Population genomics analyses of *P. polysora* from China. (A) Geographic distribution of 79 *P. polysora* isolates collected from north, central and south of China. (B) Principal component analysis of Chinese *P. polysora* population showing two genetic groups. (C) Linkage disequilibrium test of Chinese *P. polysora* population. (D) *AvrRppC* genotypes of *P. polysora* isolates in minor (red) and major (blue) genetic groups.

Due to limited samples in the minor group, we did not calculate the *F*st between two genetic groups instead of comparing the highly differentiated SNPs between two genetic clusters. A total of 32,274 SNPs (*F*st > 0.9) were filtered and these SNPs were evenly distributed in 18 chromosomes without region specificity. These variants were annotated in 6,752 protein-coding genes, of which 367 genes were affected by variants. The 20 genes were highly impacted by gaining or losing start/stop, and 250 genes showed missense variants affecting amino acid sequences. The known functional annotations of these 367 genes were mainly related to transmembrane transport, signal transduction, zinc ion/protein/nucleic acid binding, catalytic activity and protein kinase activity (Table S10). In addition, 14 of these 367 genes were annotated as secreted proteins including *AvrRppC*. The allele types of *AvrRppC* varied between the two genetic groups. In the major group, most isolates (93%) carried allele types of *AvrRppC*^1^ and *AvrRppC*^ref^. A few isolates showed the allele combination containing new allele types, *AvrRppC*^1^ to *AvrRppC*^5^ as well as *AvrRppC*^ref^ Whereas in the minor group, all four isolates carried alleles of *AvrRppC*^A^ and *AvrRppC*^J^ (Figure 7D). The CDS and amino acid sequences for new allele types are listed in Table S11.

## 4 DISCUSSION

As rust fungi are dikaryotic pathogens, obtaining their nuclear phased assembly is critical for pathogenicity studies. In this study, we reported a haplotype-phased and chromosome-scale genome of *P. polysora* based on HiFi reads and Hi-C data. The 18 chromosomes had telomeres at both ends and showed high completeness (~90% complete BUSCOs for each nucleus and 99% complete BUSCOs for two nuclei) and high continuity (N50 of scaffolds = 54 M) as well as over 80% nuclear phasing. These data strongly supported that we generated a high-quality reference genome for *P. polysora*. In addition, our study supplied a successful case to assemble a gigabase-sized fungal genome to chromosome level. The assembly pipeline we illustrated in Figure 1 will inform future genome assembly for other dikaryotic or non-haploid organisms.

### 4.1 TEs amplification leads to genome expansion of *P. polysora*, possibly in response to ancestral host decay

Compared to its close relatives, *P. polysora* has experienced a large genome expansion, with its genome size of 1.71 Gbp equivalent to 7–10 times of four other *Puccinia* species assemblies (*Pst*-104E, *Pgt*21-0, *Pt*76, and *Pca*203) (Schwessinger et al. 2018; Li et al. 2019; Wu et al. 2021; Henningsen et al. 2022). Based on a limited set of rust species, Tavares et al (2014) inferred that rusts infecting monocot Poaceae hosts have considerably smaller genomes than rusts infecting dicot Fabaceae hosts (556.6 Mbp on average). Thus, our study supplied an exception to this observation, indicating it may not be a general model. Gene duplication, repeat content variation and ploidy variation are common mechanisms to account for the genome expansion of fungi (Castanera et al. 2016; Sipos et al. 2017; Todd et al. 2017). Although gene expansion or loss has happened in the evolutionary process from *P. polysora* to other wheat-associated *Puccinia* species (Figure 6B) and correspondingly the gene number predicted in *Ppz*- GD1913 (~24000 per haplotype) is much more than those of its two close relatives, *P. coronata* (~18000) and *P. striiformis* (~14000) (Miller et al. 2018; Schwessinger et al. 2018), the expanded genes only contribute to a size difference of 7 Mbp and 17 Mbp (Table S12). On the contrary, repeat analyses suggested that genome expansion of *P. polysora* is due to its large repeat contents (85% of the total genome), notably a proliferation of LTR-RTs, which is consistent with the common evidence for rust genome expansion (Tobias et al. 2021). The evidence of genome expansion has also been detected in other obligate fungi, such as powdery mildew (PM) fungi (Spanu et al. 2010; Frantzeskakis et al. 2020) and symbiotic fungi (Miyauchi et al. 2020). PM fungi lose carbohydrate-active enzymes, transporters, etc. which are probably redundant genes in strict parasitism as tradeoffs (Spanu et al. 2010). The low abundance of PCWDEs and transcription factors in *P. polysora* as well as other *Puccinia* species was in line with the above pattern (Figure 5) (Duplessis et al. 2011, Miller et al. 2018). Despite the massive TE proliferation causing a significant enlargement in genome size, *P. polysora* shows very good gene synteny with other *Puccinia* species (Figure 5). This overall chromosome synteny was consistent with the dominant synteny detected by comparing BUSCO genes (Henningsen et al. 2022). In addition, Edwards et al (2022) reported that *Austropuccinia psidii* (Sphaerophragmiaceae, Pucciales), was in parallel with species in Pucciniaceae, Pucciales, indicating that conservatism of chromosomal synteny can exist at a family level.

In fungi, TEs have been implicated in coevolution with the host, genome architecture, plasticity, and adaptation (Lorrain et al. 2021). The “TE-thrust hypothesis” proposed that TEs can act to generate genetic novelties for organisms leading to speciation or may be related to adaptation to new challenges, e.g. new host or rapidly changing climate (Oliver and Greene 2012; Zeh et al. 2009). We detected a TE burst of *P. polysora* at around 1.7 Mya (Figure 2C) and the phylogenetic tree (Figure 6B) suggested that *P. polysora* speciated earlier than the divergence time between *P. coronata* and *P. striiformis* (40 Mya) (Aime et al. 2018). So, TE burst of *P. polysora* may be unlikely for speciation. Most scientists agree that maize originated in central Mexico and domesticated at 1–0.5 Mya from a wild grass called teosinte (Heerwaarden et al. 2011). At about 1.7 Mya, teosinte but not maize could be the major host of *P. polysora*. However, during the late Pleistocene, there may be a decline in the distribution of tropical plants, including teosinte during the cool and drying climate in Mexico (Piperno et al. 2007), leading to a drop in the effective population size of *P. polysora*. As reviewed by Lorrain et al (2021), host population dynamics can impact TEs landscapes in Fungi. The de-repressed TEs will cause genome expansion until they decay by being silenced or mutated by host defenses (Le Rouzic and Cayp 2005). Therefore, the TE burst at ~1.7 Mya could be one of the evolutionary events to support and maintain the genetic variation of *P. polysora* while going through abiotic stress and environmental change. Another explanation could be the “nearly neutral theory”, in which varied genomic features (e.g. large genome size, TE content, introns) are not initially harmful but are passively fixed by mutation or random genetic drift (Lynch and Conery 2003). With more available genome resources in the future, the relationship between effective genome size and the adaptative evolution of *P. polysora* needs to be answered.

### 4.2 Obligate biotrophic fungi lost the evolutionary feature of “two-speed genome”

The “two-speed genome” concept has been put forward to highlight the over-representation of effector-like genes in the repeat-rich and gene-sparse genome in many filamentous pathogens (Dong et al. 2015, Frantzeskakis et al. 2019). These TE-rich invasion/blocks may contribute to extensive chromosomal reshuffling and the rise of accessory chromosomes (de Jonge et al. 2013). Pathogens can resolve the evolutionary conflict through rapid evolution of effector genes to adapt to changing environments but maintain the housekeeping genes in the core genome with a moderate evolution rate (Presti et al. 2015). However, the situation is clearly different in rust fungi. There was no signal of effector compartments detected in Oat crown rust, wheat stripe rust and myrtle rust pathogens when comparing the intergenic distance between all genes and effectors (Schwessinger et al. 2018; Miller et al. 2018; Tobias et al. 2021). In *P. polysora*, the numerous TEs are widely dispersed throughout the genome (Figure 2). The intergenic distance distribution of candidate effectors overlapped with that of all genes and BUSCOs (Figure 7). This genomic architecture with TEs and genes interspersed was also reported in another biotrophic fungal group, powdery mildew (PM) fungi (Frantzeskakis et al. 2020). Similarly, AT-rich isochores or large-scale compartmentalization are also missing in PM fungi (Frantzeskakis et al. 2020). When compared with other *Puccinia* species, the high repeat content of *P. polysora* leads to much lower relative gene space (4% in *Ppz* vs 20%–35% for currently reported *Puccinia* species) but greater intergenic distance expansion (average 100 kbp of *Ppz* vs average 1–2 kbp for *Pst* and *Pca*) (Figure 4B) (Miller et al. 2018; Schwessinger et al. 2018).

### 4.3 Population genomics data accelerate the identification of potential avirulent genes

In the “arms race” between rust species and plants, the *Avr* genes in rust pathogens have been the subject of mutations to avoid the recognition by host resistance genes (Cui et al. 2015). Therefore, monitoring the variation of *Avr* genes in rust populations is a priority for disease management. Based on whole-genome SNPs, we detected two genetic groups of *P. polysora* in China. The highly differentiated SNPs between these two groups suggested mutations in 16 secreted proteins including the only verified avirulence gene, *AvrRppC*. A recent study investigated the allele types of *AvrRppC* in a Chinese *P. polysora* population (Deng et al. 2022). Isolates with allele types of *AvrRppC*^A^, *AvrRppC*^F^ and *AvrRppC*^J^ could escape the recognition of *RppC* causing southern corn rust but *AvrRppC*^ref^, *AvrRppC*^E^, and *AvrRppC*^c^ are avirulent alleles that trigger *RppC*-mediated resistance. In our study, isolates in the major group almost all carried the avirulence allele, *AvrRppC*^ref^ and four isolates in the minor genetic group carried the virulent alleles, *AvrRppC*^A^ and *AvrRppC*^J^ (Figure 7D). These results suggested that differentiation between two genetic groups could be related to virulence evolution. Similarly, this kind of relation was also detected in the wheat stripe rust pathogen (Hubbard et al. 2015). Upadhyaya et al (2021) suggested that avirulent genes of *P. graminis* showed a similar expression pattern. In this study, *AvrRppC* was classified in cluster 4 (Figure 4A) with low expression in germinated spores and early infection but high expression in intermediate infection (7 dpi). Among 14 secreted proteins with *Fst* > 0.9 between two genetic groups, FUNA_005021 (cluster 3) and FUNA_001724-T1 (cluster 5) showed similar expression patterns to *AvrRppC* and may be candidate avirulent genes. In other rust species, the rust population could drastically shift towards a wider or new spectrum of virulence over time (Miller et al. 2020; Bai et al. 2021), therefore, performing long-term surveillance as well as developing effective virulence markers are particularly critical for *P. polysora*.

### 4.4 Secretome could be an indicator to infer the presence of alternate hosts

Rust fungi have complex life cycles involving asexual as well as sexual reproduction (Figueroa et al. 2020; Zhao et al. 2021). In some species, sexual reproduction is rare or cryptic and it can remain undiscovered. For example, it took a century to discover the alternate host of *P. striiformis* (Jin et al. 2010). In our case, the sexual stage of *P. polysora* is still a mystery; teliospores are rarely found in natural fields and germination tests of teliospores were not successful in past attempts (Cammack 1959). Studies on population genetics of *P. polysora* are very limited, possibly due to the lack of effective molecular markers and the reference genome.

Our data using whole-genome SNPs supported the clonal Chinese population of *P. polysora*, however, the low *rd* between those from simulated with 0% and 50% linkage suggested the potential influence of sexual reproduction. The absence of a sexual stage or alternate host may be reflected by unequal gene numbers in the secretome. A supportive example is two species in *Melampsora*. *Melampsora lini* (autoecious, no alternate host, ~800 secreted proteins) was found to have much less secreted proteins and secreted plant cell wall-degrading enzymes (PCWDEs) than its close relative, *M. larici-populina* (heteroecious, with both primary and alternate hosts, ~1800 secreted proteins) (Presti et al. 2015). In our case, we identified 2,434 secreted proteins for *P. polysora*, which was comparable to the secretomes of two heteroecious relatives, *P. coronata* (~2500) and *P. graminis* (*~*2500) (Miller et al. 2018; Li et al. 2019). Of course, the single *Melampsora* example did not allow for making a general conclusion. Varied quality of genome assemblies and different prediction methods can lead to significant bias in secreted protein predictions. However, it will be valuable to test this hypothesis with more available high-quality rust genomes in future studies. If the hypothesis is true, our results may suggest the existence of alternate hosts that have not been discovered yet for *P. polysora*.

## 5 CONCLUSION

In conclusion, we successfully obtained the first nuclear phased and chromosome-scale genome assembly of the serious fungal pathogen, *P. polysora*. The high-quality genome assembly and fine-time-point transcriptome facilitate the comparative genomic analyses. The genome expansion of *P. polysora* is driven by TEs, but this did not affect its gene synteny with close relatives. TEs insertion in turn led to expanded intergenic distance. The investigation of genome characteristics and functional features provided a fundamental resource for pathogenicity studies as well as to understand the evolutionary mechanisms of genome expansion in rust fungi. In addition, the population genomic data revealed low genetic differentiation in the Chinese *P. polysora* population, which expanded from south to north in recent decades. However, a minor genetic group with low frequency suggested the ongoing virulence evolution to evade recognition by *RppC*, a major resistance gene in Chinese corn cultivars, alarming the need to excavate additional resistance genes as soon as possible.

## ACKNOWLEDGEMENTS

We would like to thank Xiufeng Liu, Peng Lu, Gongxian Liao, Shikun Pan, Xingshang Lu, Baoqin Tan, Qiusheng Luo for field diseased sample collection. We also appreciate valuable suggestions from Dr. Mao Jianfeng (Beijing Forestry University, Beijing) and Dr. Qi Wu (Institute of Microbiology, Chinese Academy of Sciences, Beijing) before submission. Clement K.M. Tsui is grateful to CAS PIFI for the award of visiting scientist fellowship.

## AUTHOR CONTRIBUTIONS

The experiment was conceived and managed by Lei Cai and Junmin Liang. Field samples were collected by Junmin Liang, Yuanjie Li, Zhanhong Ma, Leifu Li and Keyu Zhang. Single-spore isolation, reproduction, DNA/RNA extraction and other biological materials used for sequencing were conducted by Junmin Liang and Yuanjie Li. Data analyses were performed by Junmin Liang. Analysis tools were assisted by Peter N. Dodds, Melania Figueroa and Jana Sperschneider. Junmin Liang wrote the manuscript and Peter N. Dodds, Jana Sperschneider, Clement K.M. Tsui, and Lei Cai reviewed the manuscript.

### DATA AVAILABILITY

All raw sequence reads (PacBio, Illumina, Hi-C and RNA-seq data) generated in this study are available both in NCBI and the National Microbiology Data Center (NMDC, https://nmdc.cn/) under the BioProject of PRJNA881038 and NMDC10018113, respectively. The accession number for all raw data are listed in Table S13. The accession number for assembled genome was JAOOQA000000000 in NCBI and NMDC60042795 in NMDC. Scripts and pipelines used to construct figures and annotation files (gff3 and cds) are available at https://github.com/jimie0311/Puccinia-polysora-genome.

### FUNDING

This work was financially supported by the National Sciences Foundation of China (NSFC 31972210 and NSFC 31725001) and the National Sciences and Technology Fundamental Resources Investigation Program of China (2021FY100900).

## Supplementary files

Table S1 Information of 79 isolates from China used for genome resequencing.

Table S2 Contig information removed from the final assembly due to low-quality assessment.

Table S3 Treatments for duplicate contigs between/within haplotype A and haplotype B

Table S4 Sample information and data used for transcript analyses.

Table S5 Functional annotation of specific orthologs between two haplotypes.

Table S6 GO and IPR annotation for predicted candidate effectors.

Table S7 Features of secreted proteins in different expression clusters of *P. polysora* GD1913.

Table S8 Proteome comparison of four *Puccinia* species.

Table S9 GO annotation of top 42 *P. polysora* specific orthogroups with high gene copy number. Table S10 Table S10 Variants impact of SNPs located in 367 genes with *F*st > 0.9 between two groups.

Table S11 The CDS and amino acid sequences of *AvrRppC* gene.

Table S12 Comparison of Gene number and gene length among three *Puccinia* species.

Table S13 NMDC numbers for sequencing raw data used in this study.

Figure S1 Genome size estimation of *Ppz*-GD1913 using 90 Gb illumina short reads.

Figure S2 GO enrichment analysis of secreted proteins on two haplotypes.

Figure S3 Pair-wise synteny of *P. polysora* and three close relatives.

Figure S4 The elbow bend plot.

Figure S5 The bi-allele distribution of 79 isolates used for population genomics. Two isolates (GD1922-3 and GX1905-2) in red squares are removed from the final analysis due to abnormal distribution.

