## Supplementary figures and images for "Haplotype-phased and chromosome-level genome assembly of *Puccinia polysora*, a giga-scale fungal pathogen causing southern corn rust"

### Figure S1. Genome size estimation.pdf

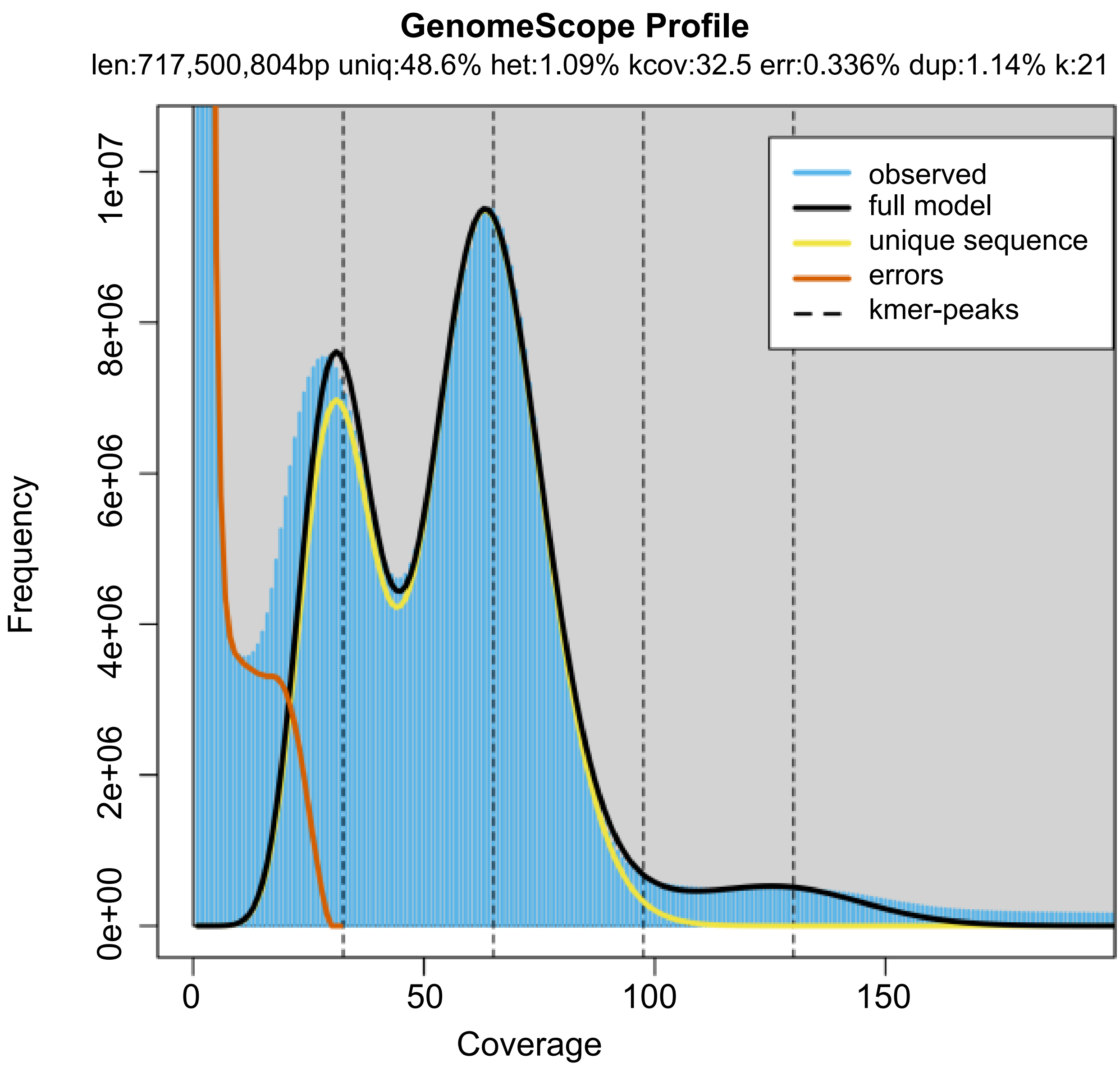

### Figure S2 GO enrichment of secreted proteins on two haplotypes.pdf

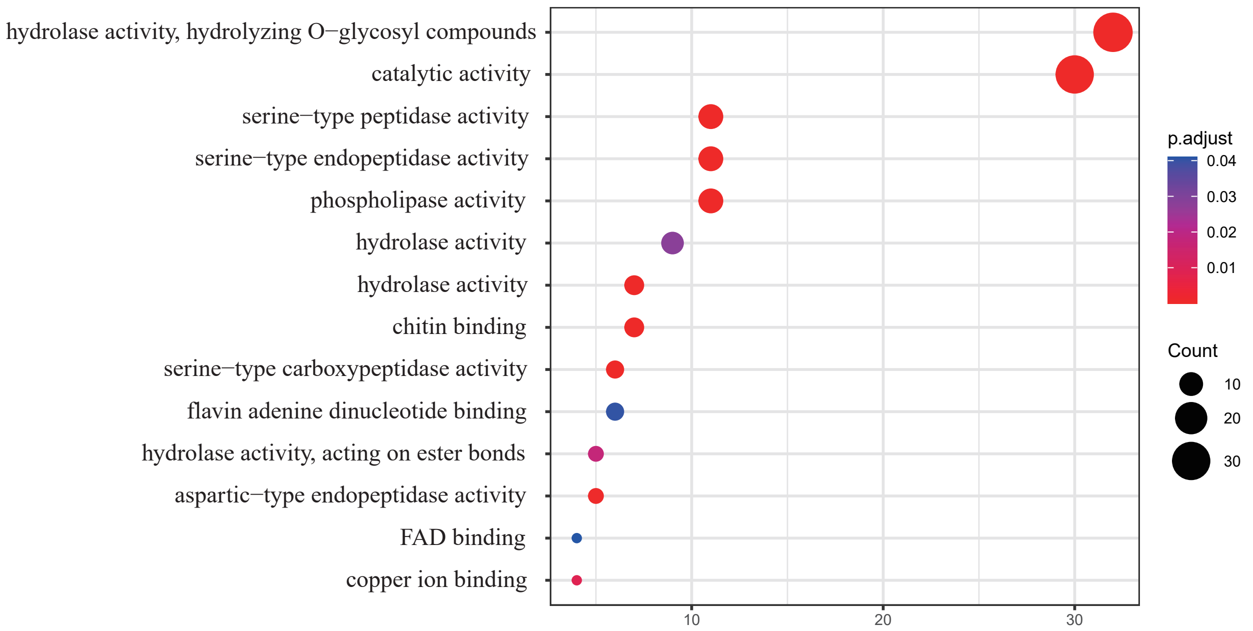

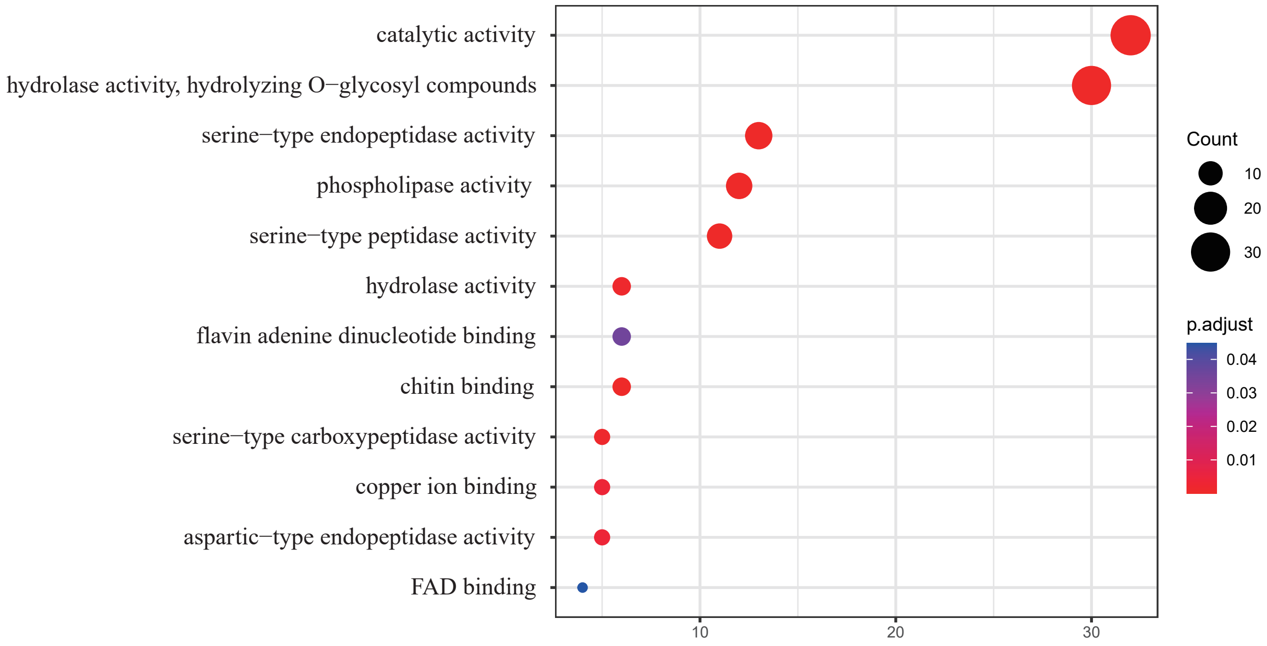

### Figure S3 macro-syndeny.pdf

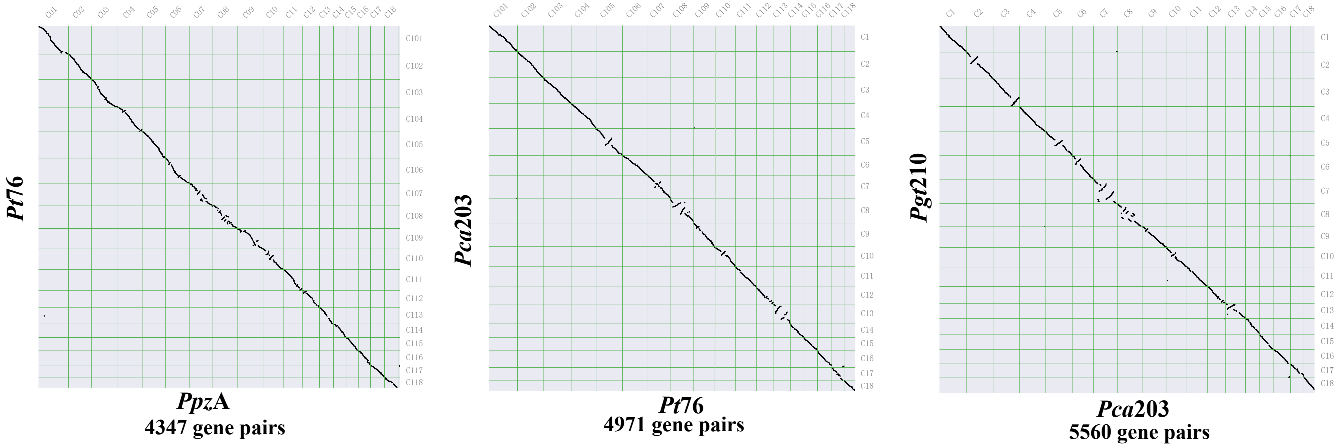

### Figure S4 The elbow bend.pdf

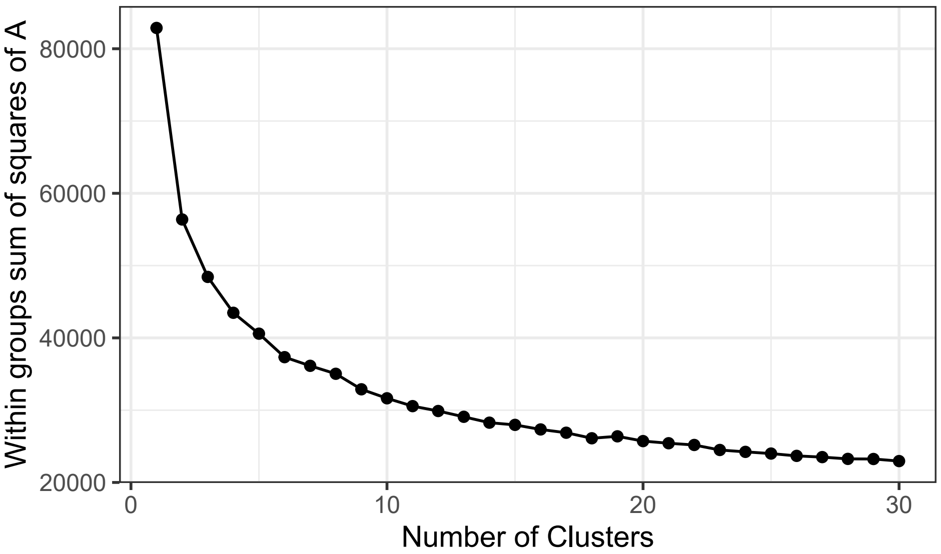

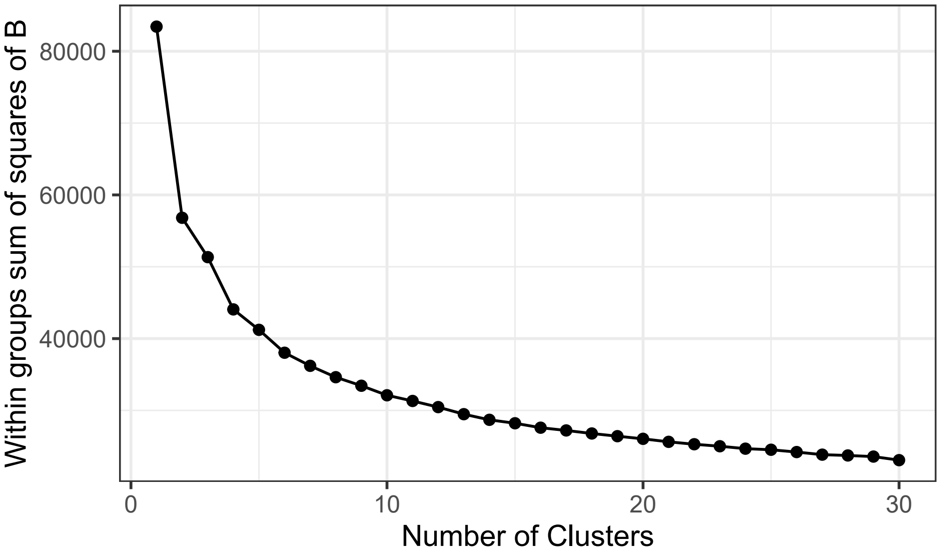

### Figure S5 allele balance.pdf

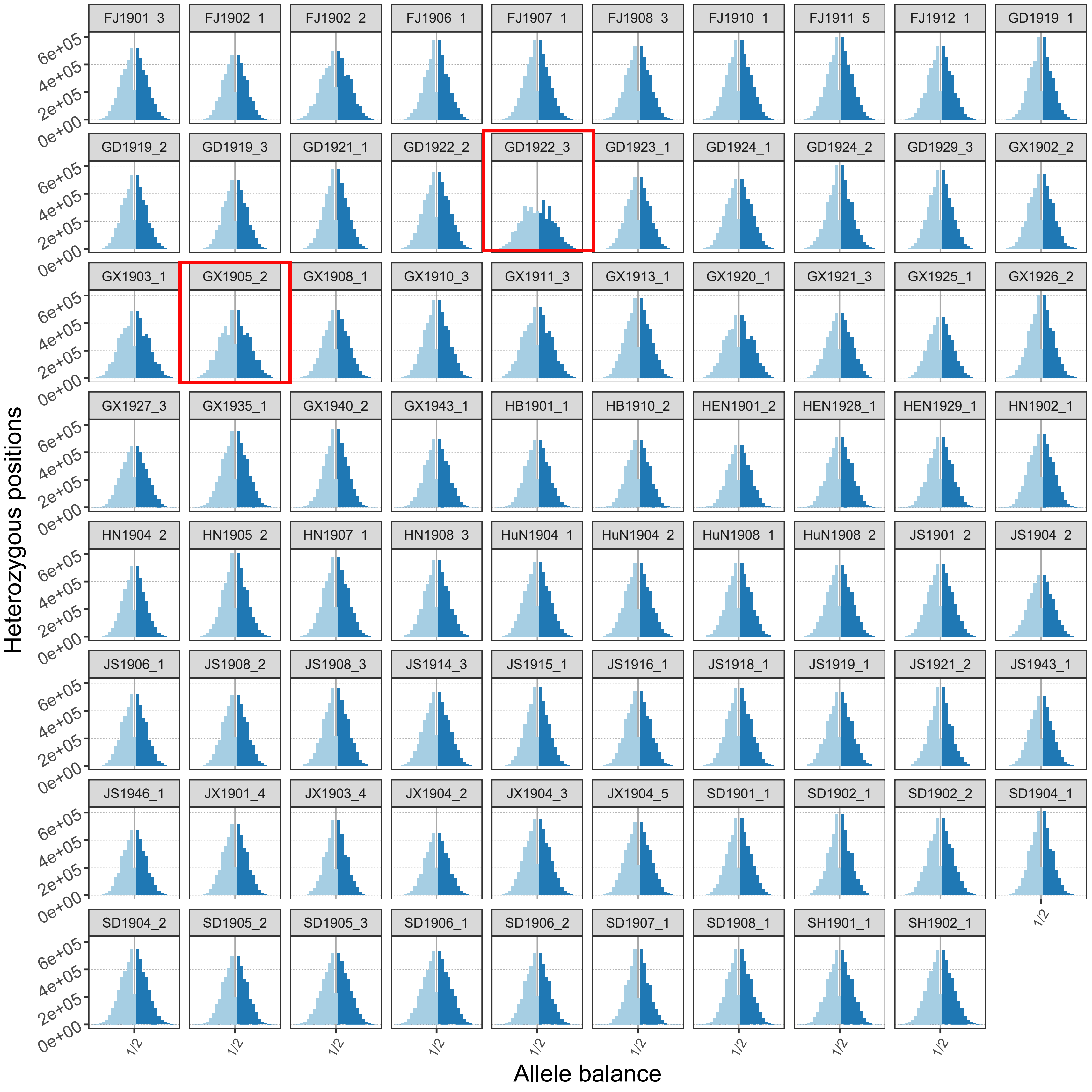
